# Multimodal characterization of variation in neuronal types in the mouse basal ganglia

**DOI:** 10.64898/2026.06.24.734325

**Authors:** Agata Budzillo, Matt Mallory, Rachel Dalley, Changkyu Lee, Xuehan Ci, Xiao-Ping Liu, Sarah Walling-Bell, Rusty Mann, Nelson Johansen, Ellie Adams, Lauren Alfiler, Julia Andrade, Angela Ayala, Katherine Baker, Stuard Barta, Zoe Benaissa, Darren Bertagnolli, Ashwin Bhandiwad, Krista Blake, Phil Bohn, Krissy Brouner, Trangthanh Cardenas, Tamara Casper, Scott Daniel, Nadezhda I. Dotson, Tom Egdorf, Rachel Enstrom, Amanda Gary, Jeff Goldy, Kristen Hadley, Zoe C. Juneau, Megan Koch, Gabriela Leon, Jocelin Malone, Arena Manning, Rachel McCue, Audrey McCutcheon, Medea McGraw, Kamiliam Nasirova, Lindsay Ng, Alana Oyama, Christina Alice Pom, Lydia Potekhina, Ramkumar Rajanbabu, Shea T. Ransford, Ingrid Redford, Christine Rimorin, Kara Ronellenfitch, Augustin Ruiz, Michael Tieu, Jessica Trinh, Sara Vargas, Maria Camila Vergara, Jeanelle Ariza, Nick Dee, Luke Esposito, Kathryn Gudsnuk, Samantha Delmont Hastings, Lauren Kruse, Kimberly A. Smith, Susan M. Sunkin, Quanxin Wang, Jack Waters, Ed S. Lein, Jonathan T. Ting, Bosiljka Tasic, Zizhen Yao, Hongkui Zeng, Brian Kalmbach, Tim Jarsky, Staci A. Sorensen, Brian R. Lee, Nathan W. Gouwens

## Abstract

The basal ganglia (BG) are a set of topographically organized, interconnected structures that are pivotal for regulating volitional movement and other aspects of cognitive, motivational, and affective behavior. Recently generated taxonomies of transcriptomically-defined cell types (T-types) have revealed both fine-grained distinctions in gene expression between neurons in these structures as well as continuous transcriptomic variation across similar T-types^1–9^, which are both related to location within a structure. However, it remains unclear to what extent these and other cellular properties co-vary with each other. Therefore, we performed Patch-seq experiments^10,11^ on over 900 neurons in mouse brain slices from BG to provide an integrated view of the co-variation between gene expression, location, physiology, and morphology measured from the same neurons. Medium spiny neurons (MSNs) from both the direct and indirect pathways across the dorsal and ventral striatum follow a gene expression gradient that varies in a dorsolateral to ventromedial direction; we find that this gradient also corresponds with systematic differences in action potential kinetics and dendritic arborization. Our analysis also characterizes additional multimodal dimensions of MSN variation, such as those between direct and indirect pathway neurons and between matrix and striosome neurons. Furthermore, through comparison with Patch-seq data from macaque, we demonstrate that the relationship between the transcriptomic/spatial gradient and electrophysiological and morphological properties is conserved across these two species. We also find that properties of striatal interneurons, such as action potential kinetics, vary across the striatum in a manner consistent with the MSN gradient. Outside the striatum, our multimodal Patch-seq dataset from the globus pallidus, subthalamic nucleus, and substantia nigra enabled us to characterize transcriptomically-defined types and link them to prior descriptions of cell types in these structures. Finally, we examined to what extent the MSN transcriptomic/spatial gradient persisted across different stages of the BG circuit by comparing Patch-seq neurons to reconstructed whole-neuron morphologies and the topography of their interareal projections, finding that the gradient is better preserved in GPe and GPi compared to SNr. Our study links transcriptomic variation across T-types in mouse BG to spatial localization and phenotypic differences at the level of individual cells, improving our understanding of cell type architecture in topographically organized circuits of the brain.

## Introduction

The basal ganglia are a set of interconnected nuclei that regulate action selection and motor control^12^. A simplified but highly influential model of basal ganglia organization divides the circuit into direct and indirect pathways, arising from the recruitment of *Drd1* or *Drd2* receptor-expressing medium spiny neurons (MSNs) in the striatum, connecting to the thalamus either directly via the substantia nigra pars reticulata (SNr) and globus pallidus internal segment (GPi) to promote movement, or indirectly, with a sign-reversing stopover in globus pallidus external segment (GPe), to inhibit movement^13–15^. In addition to this canonical pathway structure, there is substantial evidence that basal ganglia circuits are organized topographically in parallel input–output streams^16–20^. At a high level, this is reflected in the preferential targeting of sensorimotor and associative cortical and thalamic inputs to distinct regions of the caudoputamen (CP), and limbic inputs to the nucleus accumbens (ACB). The topography of cortical and thalamic inputs to striatum, as well as evidence of spatial structure in striatum^3,4,6,21^ and other BG nuclei^22,23^, suggests that location within these structures is linked to cell type identity.

Considerable progress has been made in describing the diverse cellular landscape within BG structures in rodents and primates through classification of their molecular, phenotypic, and connectivity profiles. In the past decade, single cell RNA sequencing has expanded the landscape of basal ganglia cell types, uncovering subtleties in molecular diversity within each region^1–9^. Despite these advances, establishing how molecular diversity relates to phenotypic, spatial and functional variation across basal ganglia structures remains a challenge; however, identifying those connections could enable the development and use of genetic tools to access specific cell type populations with distinct functional properties, which could benefit both experimental and therapeutic applications.

To establish these links, we collected transcriptomic and electrophysiological data from 904 cells along with reconstructed morphologies from 192 of those cells across multiple basal ganglia regions in the mouse using the Patch-seq technique. Recorded cells were registered within the Allen Common Coordinate Framework (CCFv3)^24^ to assess spatial differences in multimodal properties. This multimodal approach allowed us to link traditional cell classifications to discrete types in a well-defined whole brain molecular taxonomy^2^, while also investigating continuous co-variation in properties. Using these data, which we believe represent the most comprehensive multimodal dataset from the basal ganglia to date, we were able to identify specific gene expression patterns that were predictive of electrophysiological and morphological features and characterize specific transcriptomically-defined cell types. We also integrated the Patch-seq dataset with a previously published set of MSN whole-neuron morphologies to investigate how the systematic multimodal variation we observed translated to projection patterns in downstream basal ganglia structures.

## Results

### Collection of Patch-seq data in basal ganglia

In order to study the correspondence between gene expression, physiological, morphological properties and spatial location in individual basal ganglia neurons, we performed Patch-seq experiments in acute brain slices from adult mice containing CP, ACB, OT, GPe, GPi, subthalamic nucleus (STN), SNr, ventral tegmental area (VTA) or substantia nigra pars compacta (SNc), using a combination of wild-type mice, transgenic mice, and viral labeling tools to target diverse cell types (Fig. 1a; Supplementary Data 1-2). We collected responses to a standardized set of hyperpolarizing and depolarizing current injections and, in a subset of cells, spontaneous firing activity. During the recording, cells were filled with biocytin for later staining to perform post-hoc morphological reconstruction and mapping of spatial location to the Allen Common Coordinate Framework (CCFv3)^24^, Fig. 1b). After recording, we aspirated the nucleus and cytosol of the cell for single-cell RNA sequencing. This study includes 1,296 Patch-seq experiments with high-quality transcriptomic (T) data collected from 488 mice. Among these, 904 experiments also yielded high-quality electrophysiological (E) recordings, while dendritic morphologies (M) were reconstructed for 260 neurons, including 192 with triple modality (MET). Local axons were reconstructed when visible for interneuron populations. In our dataset, 702 neurons were acquired from striatal regions (dorsal striatum or CP, and ventral striatum including ACB and OT, Fig. 1b). The remaining 152 neurons were acquired from GPe (35), GPi (19), STN (40), SNr (42), and VTA / SNc (16).

**Figure 1:**
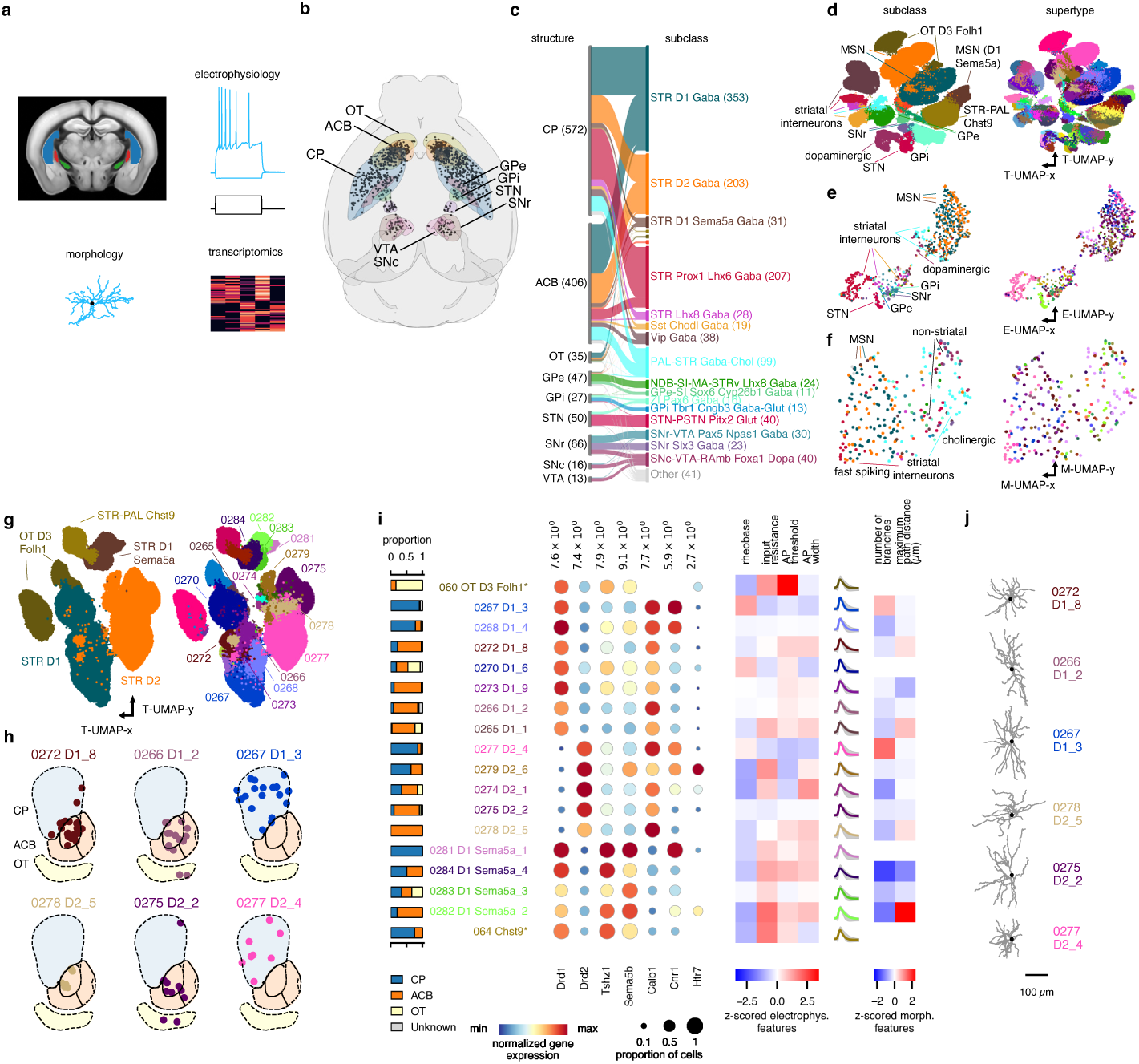
Overview of experimental strategy, basal ganglia types and diversity of MSN types. **a**, Schematic of the Patch-seq method. Single cells were targeted in coronal slices of CP (blue), GPe (red), GPi (green), ACB, OT, SNc, SNr, STN and VTA (not shown) for whole-cell current clamp recording (electrophysiology), reconstruction (morphology), and gene expression analysis (transcriptomics). **b**, Somata from Patch-seq experiments registered to the Allen CCFv3 (n=1,156). **c**, River plot showing the relationships between BG structure and transcriptomic subclass for Patch-seq experiments. **d**, UMAP of gene expression of reference dissociated cells (n=159,538) and Patch-seq cells (1,296). (MSN: STR D1 Gaba, STR D2 Gaba, STR D1 Sema5a Gaba; striatal interneurons: STR Prox1 Lhx8 Gaba, STR Lhx8 Gaba, Sst Chodl Gaba, Vip Gaba, PAL-STR Gaba-Chol; GPi: ZI Pax6 Gaba, GPi Tbr1 Cngb3 Gaba-Glut; GPe: NDB-SI-MA-STRv Lhx8 Gaba, GPe-SI Sox6 Cyp26b1 Gaba; STN: STN-PSTN Pitx2 Glut; SNr: SNr-VTA Pax5 Npas1 Gaba, SNr Six3 Gaba; dopaminergic: SNc-VTA-RAmb Foxa1 Dopa). UMAPs in *d-g* are colored by subclass (left) and supertype (right). **e**, UMAP of electrophysiology sparse PCs (n=904 Patch-seq cells). **f**, UMAP of dendritic morphological features (n=260 Patch-seq cells). **g**, UMAP of gene expression of reference dissociated cells (n=95,625) and Patch-seq cells (n=624) in MSN (STR D1 Gaba, STR D2 Gaba, STR D1 Sema5a Gaba) and MSN-adjacent (OT D3 Folh1 Gaba, STR-PAL Chst9 Gaba) subclasses, colored by subclass (left) and supertype (right). ”Gaba” has been removed from subclass labels for brevity. **h**, Coronal sections of Allen CCFv3 showing registered Patch-seq MSN soma locations (colored by supertype). ”STR” and ”Gaba” have been removed from MSN supertype labels for brevity. **i**, Summary of multimodal features for select MSN and MSN-adjacent supertypes. From left: proportion of Patch-seq somas registered to CP/ACB/OT; select marker gene expression, normalized to maximum expression of each gene across MSNs; select z-scored electrophysiology features; average action potential waveform and its variability (±2 standard deviations; all MSN average in gray); select z-scored morphology features. **j**, Example dendritic reconstructions for select MSN supertypes.

To link cellular phenotypes with transcriptomic identity, we mapped each cell to a reference whole mouse brain taxonomy^2^ composed of several increasingly granular hierarchical levels: class, subclass, supertype, and cluster (aka transcriptomic type, or T-type) (Methods). Each Patch-seq cell’s transcriptome was mapped to the finest resolution (T-type); however, certain analyses were performed while grouping cells at the subclass or supertype level. Transcriptomic subclass assignments for the dataset can be seen in Figure 1c. To visualize transcriptomic, electrophysiological and morphological diversity, we applied Uniform Manifold Approximation and Projection (UMAP)^25^ to the multidimensional feature spaces for each modality (Fig. 1d-f). Cells mapping to different transcriptomic subclasses were typically found in distinct locations in both transcriptomic and electrophysiology feature spaces (Fig. 1d-e, left); in many cases this was true for supertypes (Fig. 1d-e, right) and T-types, as well (Extended Data Fig. 2; Extended Data Fig. 3). Although the dendritic feature space was less structured compared to the other two modalities, MSNs and fast spiking striatal interneurons formed a separate island from cholinergic striatal interneurons and other types of basal ganglia neurons (Fig. 1f; Extended Data Fig. 4).

### Multimodal characterization of MSN cell types

Striatal MSNs, the principal projection neurons of the striatum, exhibit substantial molecular and functional diversity^3–5,26^. In the reference taxonomy^2^, MSNs are distributed across 20 transcriptomic supertypes within three subclasses: STR D1 Gaba (D1), STR D2 Gaba (D2), STR D1 Sema5a Gaba (D1 Sema5a) (Fig. 1g). D1 and D2 MSNs express *Drd1* or *Drd2* and contribute to the direct and indirect output pathways, respectively, whereas D1 Sema5a corresponds to MSNs with a transcriptomic profile distinct from both D1 and D2 and scattered distribution across the striatum^27^. This subclass also contains a supertype (281 STR D1 Sema5a Gaba 1) that includes a subset of cells expressing both *Drd1* and *Drd2*^27^. We collected a small number of cells from two additional transcriptomically related subclasses: OT D3 Folh1 Gaba (D3 Folh1), which defines small *Drd3+* GABAergic neurons in the Islands of Calleja of the OT, and STR-PAL Chst9 Gaba (Chst9), which defines cells that are transcriptomically similar to STR D1 Sema5a and are found in scattered locations and streaks in the striatum and neighboring structures^27^.

Patch-seq cells mapping to different MSN supertypes had varied distributions within striatum (Fig. 1h, i left), consistent with spatial transcriptomic evidence^27^. Molecular diversity was also associated with phenotypic differences across MSN supertypes and MSN-adjacent cells; for example, action potential (AP) width varied across MSNs (median = 0.84 ms; IQR = 0.94 ms–0.74 ms; range = 1.98 ms–0.3 ms; *n* = 450), with some of the fastest APs observed in cells mapping to dorsolaterally-located supertypes 0267 STR D1 Gaba 3 (0.75 ms ± 0.14 ms, *n* = 65) and 0277 STR D2 Gaba 4 (0.75 ms ± 0.11 ms, *n* = 46) and wider APs observed in ventromedially-located supertypes 0272 STR D1 Gaba 8 (0.94 ms ± 0.20 ms, *n* = 64) and 0278 STR D2 Gaba 5 (0.95 ms ± 0.15 ms, *n* = 9). Input resistance varied across groups, with the highest values seen in cells mapping to D3 Folh1, D1 Sema5a, and Chst9 subclasses, as well as the D2 supertype 0279 STR D2 Gaba 6 (Fig. 1i). Dendritic arbors also varied considerably across supertypes, with highly complex dendrites and a high number of branches in dorsolaterally-biased supertypes 0267 STR D1 Gaba 3 (50.86 ± 10.89, *n* = 14) and 0277 STR D2 Gaba 4 (59.92 ± 21.77, *n* = 12), and sparser dendritic arbors with fewer branches in ventromedial supertypes 0266 STR D1 Gaba 2 (27.80 ± 10.43, *n* = 10) and 0275 STR D2 Gaba 2 (34.00 ± 10.77, *n* = 18). Together, these results point to correspondences between transcriptomic diversity, location within the striatum, and intrinsic electrophysiological and morphological variation in MSNs.

We used the cells in our Patch-seq dataset that contained all three data modalities (morphology, electrophysiology, and transcriptomics) to define MET-types as done previously for mouse neurons in visual cortex^28,29^ (Extended Data Fig. 1). Interestingly, while we could identify clearly defined MET-types for several sets of cells in GPe, GPi, STN, SNr, and SNc/VTA, as well as sets of striatal interneurons (see below), we found that the MSN population was represented by multiple closely related MET-types (Extended Data Fig. 1b). Unlike most non-MSNs, MSN T-types were typically split and re-merged across multiple MET-types. This suggested that MET-type definitions may be less robust for MSNs than for other neuronal populations and that alternative analyses that characterize continuous variation across multiple modalities could be more informative for these cells.

### Continuous multimodal co-variation in MSNs

Therefore, to characterize the relationship between the transcriptomic landscape and cellular phenotypes in MSNs more directly, we analyzed the MSN Patch-seq dataset using sparse reduced-rank regression (sparse RRR)^30^, a method that identifies gene expression patterns most predictive of electrophysiological and morphological features by defining a lower-dimensional latent space and sparse set of contributing genes. We applied sparse RRR to the Patch-seq dataset of D1, D2, and D1 Sema5a MSNs separately for electrophysiology and dendritic morphology features, yielding sets of both electrophysiology (E-LF) and morphology (M-LF) latent factors (Fig. 2a, p). The number of latent factors and other hyperparameters were optimized using cross-validation (Extended Data Fig. 5; Methods). Relationships between genes and features within the latent factor spaces are visualized with bi-plots (Fig. 2b, q; Extended Data Fig. 6), in which LFs estimated from gene expression and features are shown side-by-side and annotated with the most strongly correlated genes and features.

**Figure 2:**
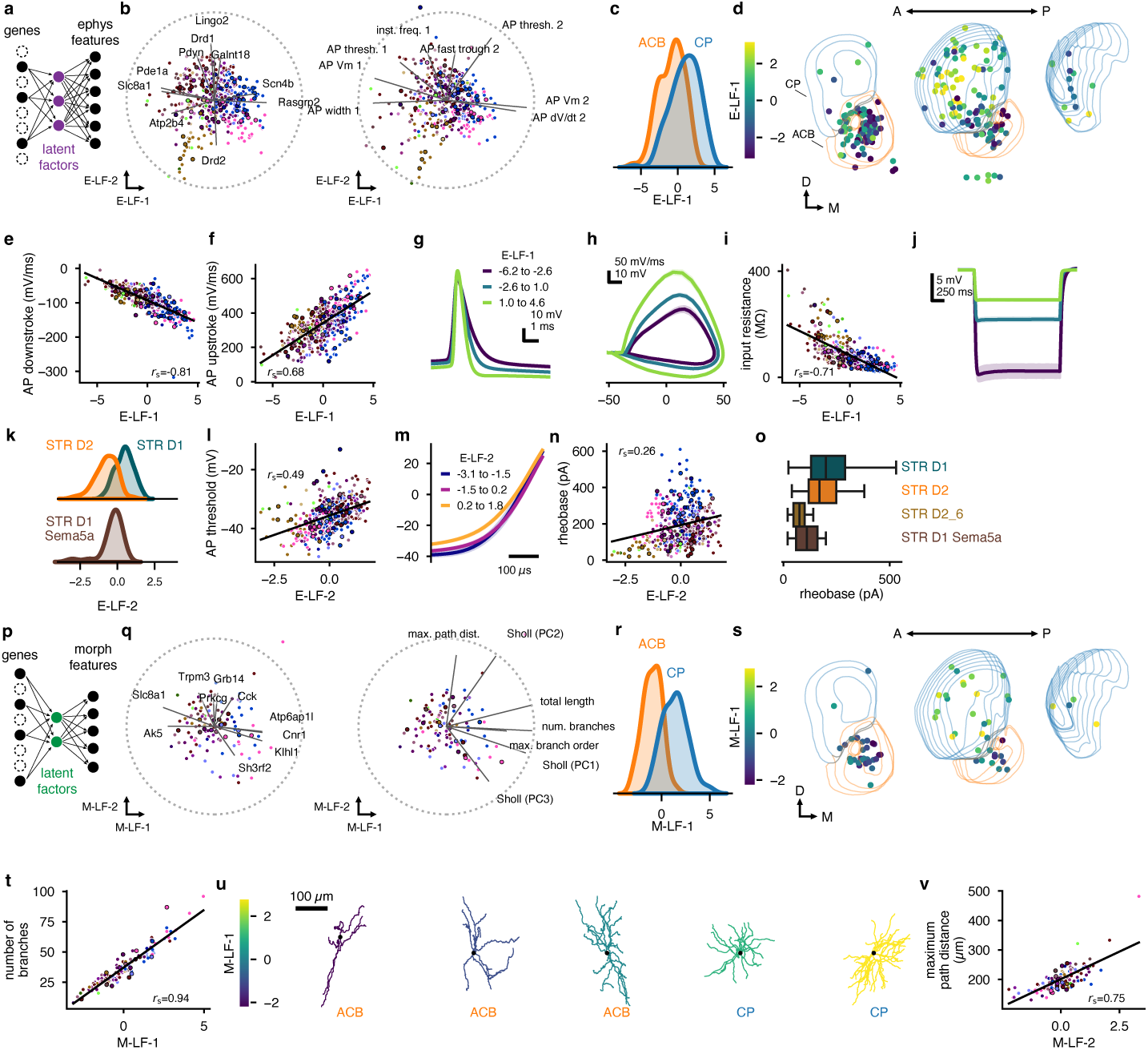
Continuous multimodal co-variation in MSNs. **a**, Schematic of sparse reduced-rank regression (sparse RRR) of electrophysiology (ephys) features in MSNs. Latent factor (LF) values are derived from a weighted subset of differentially-expressed genes and used to predict ephys features. **b**, Bi-plots of E-LF-1 and E-LF-2 for 450 MSNs. Genes (left) and ephys features (right) with the highest LF correlations are shown. A correlation value of one is indicated by the dotted circle. Color indicates supertype, as labeled in Fig. 1; black borders indicate held-out cells not used to fit the regressions. **c**, Kernel density estimates (KDEs) of E-LF-1 distribution in MSNs acquired from ACB (n=228) or CP (n=176). **d**, Pseudo 3D plots of successive sections of CP, ACB, and OT. MSN soma locations are colored by E-LF-1 value. **e,f,i**, Relationships between E-LF-1 and ephys features; lines are regression fits. Cells are colored by transcriptomic supertype. **g,h,j**, Averaged AP waveforms (**g**), AP phase plots (**h**), and subthreshold voltage responses (**j**) for MSNs grouped by E-LF-1 value. **k**, KDEs of E-LF-2 distribution in D1 (n=269), D2 (n=156), and D1 Sema5a (n=25) MSN subclasses. **l,n**, Relationships between E-LF-2 and ephys features; lines are regression fits. **m**, Averaged AP rising phase for cells grouped by E-LF-2 value. **o**, Box plots highlighting low rheobase current in 0279 STR D2 Gaba 6 supertype compared to other MSN subclasses. ”Gaba” has been removed from subclass labels for brevity. **p**, Same as (**a**) but for morphological (morph) features. **q**, Bi-plots of M-LF-1 and M-LF-2 for 117 MSNs. Similar to (**b**), genes and morph features with the highest LF correlations are shown, and cells are colored by supertype. A correlation value of one is indicated by the dotted circle. **r**, KDEs of M-LF-1 distribution in MSNs derived from ACB (n=67) or CP (N=49). **s**, Same as (**d**); MSN soma locations are colored by M-LF-1 value. **t,v**, Relationship between M-LF-1 and the number of dendritic branches. **u**, Example dendritic MSN morphologies, ordered and colored by M-LF-1 value. **v**, Relationship between M-LF-2 and the maximum dendritic path distance.

The first electrophysiology latent factor (E-LF-1) was associated both with categorical location within the striatum (CP versus ACB), and in a more continuous fashion, exhibiting low values at the ventromedial end of the striatum and higher values toward the dorsolateral end (Fig. 2c-d). This transcriptomic gradient described by E-LF-1 was highly associated with AP shape and kinetics, as well as with input resistance (Fig. 2b, e-j). The most strongly contributing genes to E-LF-1 included voltage-gated sodium channel *β* auxiliary subunit *Scn4b*, which regulates sodium channel gating and action potential dynamics^31^, Rap1 guanine nucleotide exchange factor *Rasgrp2*, which has been linked to dopamine-dependent modulation of MSN excitability^32^, and sodium-calcium exchanger *Slc8a1*. MSNs with low E-LF-1 values had shallower AP rising and falling phases and higher input resistances than those with high E-LF-1 values. This association appeared continuous throughout the striatum and was shared by both D1 and D2 MSNs.

The second electrophysiology latent factor (E-LF-2) was driven most strongly by *Pdyn*, *Drd1*, *Lingo2*, and *Drd2* and distinguished between D1 and D2 MSNs (Fig. 2k). Unlike E-LF-1, E-LF-2 did not have a strong spatial bias, but was associated with the excitability of cells given its correlation with AP threshold and rheobase (Fig. 2k-n). Notably, cells mapping to the 0279 STR D2 Gaba 6 supertype had particularly low values of E-LF-2 (Fig. 2b, brown) and electrophysiological features outside of the typical D2 MSN range (Fig. 2l,n), suggesting that these cells are among the most excitable of MSNs. Rheobase current varied significantly across the four groups examined (one-way ANOVA, F(3, 446) = 25.20, *p <* 0.001): D1, D2 excluding the 0279 STR D2 Gaba 6 supertype, D1 Sema5a, and 0279 STR D2 Gaba 6; significant differences were seen between all pairs except 0279 STR D2 Gaba 6 and D1 Sema5 Gaba (post-hoc Tukey tests; Fig. 2o).

Sparse RRR was similarly predictive for both electrophysiology and morphology features in MSNs (electrophysiology: CV-*R*^2^ = 0.198, morphology: CV-*R*^2^ = 0.221). M-LF-1 was highly correlated with E-LF-1 (Extended Data Fig. 5d). The top contributing genes to this transcriptomic gradient were the cannabinoid receptor *Cnr1*, which has previously been described as varying continuously across this axis of the striatum^3,4,21^, *Klhl1*, *Slc8a1*, and *Ak5*. M-LF-1 had a strong association with the ventromedial to dorsolateral axis in striatum (Fig. 2r-s). Dendritic arbor complexity varied systematically along M-LF-1, with sparser, less complex dendritic arbors at low M-LF-1 values (found in ventromedial MSNs) and more complex arbors with a greater number of dendritic branches at high M-LF-1 values (found in dorsolateral MSNs) (Fig. 2t-u). A second morphology latent factor, M-LF-2 was associated with the maximum length of individual dendritic branches (Fig. 2q, v).

### Additional variation in MSN cell types

The electrophysiological and morphological latent factors account for a substantial fraction of variation across the MSN population, relating to both location and membership in direct or indirect pathways. Another well-established dimension of MSN diversity is whether they reside in patches (striosomes) or the matrix: neurochemically distinct compartments which are thought to contribute differentially to limbic and sensorimotor learning^33^. Several MSN supertypes have spatial patterns largely consistent with either patch or matrix compartmentalization^27^ (Fig. 3a). In MERFISH data, 0267 STR D1 Gaba 3 cells appear enriched in matrix, while 0268 STR D1 Gaba 4 have a more patch-like expression pattern. For D2 MSNs, supertype 0277 STR D2 Gaba 4 includes T-types that appear predominantly matrix-enriched, with the exception of T-type 0979, which has an expression pattern more consistent with patches (Fig. 3a). Another transcriptomic supertype, 0279 STR D2 Gaba 6, also has a patch-like spatial pattern. We examined the gene expression of Patch-seq cells assigned to these four representative matrix-like and patch-like MSN supertypes, while also analyzing T-type 0979 separately from other 0277 STR D2 Gaba 4 cells, and confirmed expression of marker genes associated with matrix (*Rasgrp2, Id4, Calb1*) or patch (*Sema5b, Oprm1*) identity (Fig. 3b).

**Figure 3:**
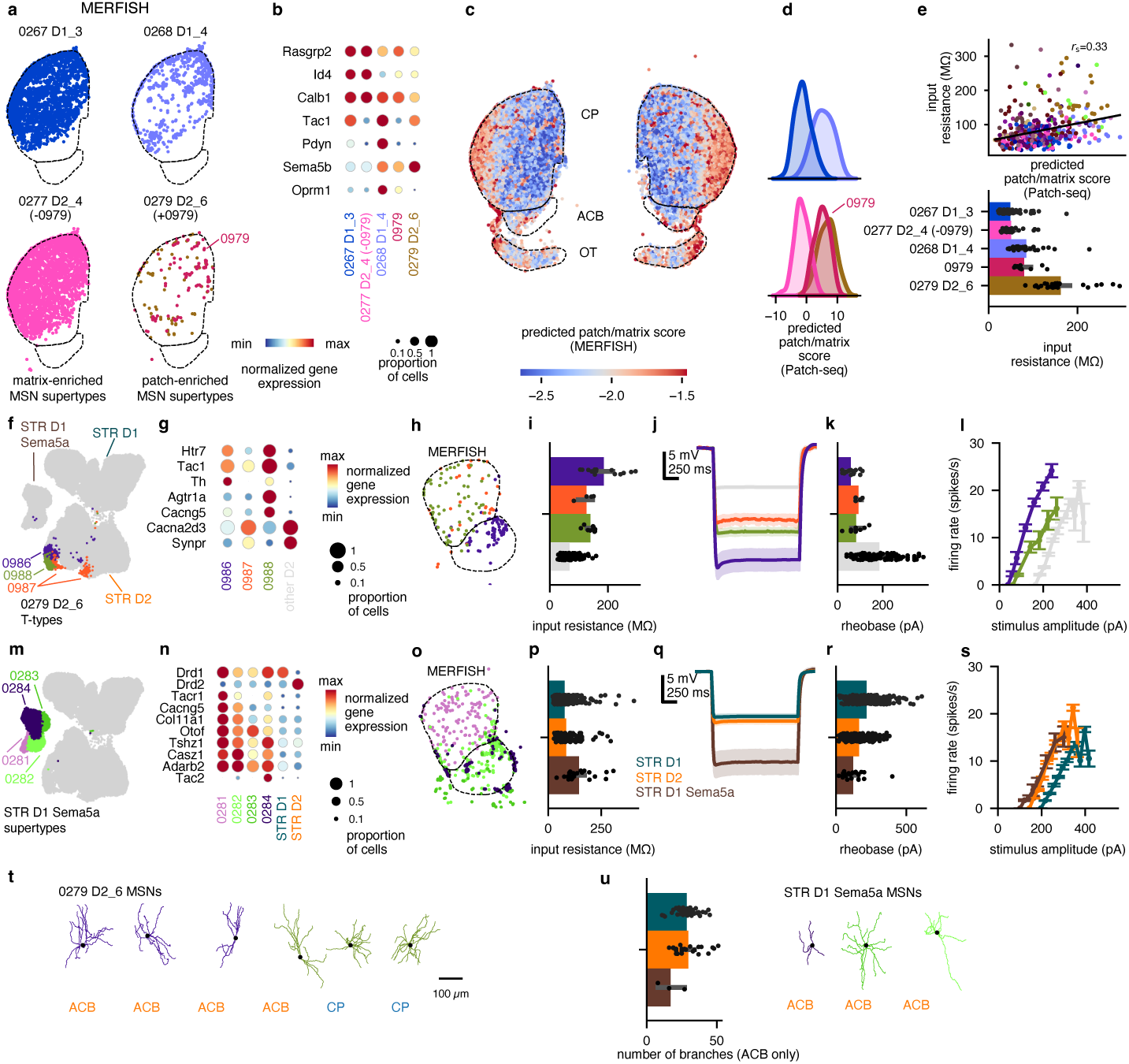
Characterization of additional dimensions of multimodal variation in MSNs. **a**, Representative MERFISH section of the adult mouse brain containing CP and ACB, colored by matrix-enriched (left) patch-enriched (right) supertypes. T-type 0979 STR D2 Gaba 4 has been analyzed separately from the otherwise matrix-enriched supertype 0277 STR D2 Gaba 4 and included in the 0279 STR D2 Gaba 6 panel. ”STR” and ”Gaba” have been removed from supertype labels in figure panels for brevity. **b**, Marker gene expression in Patch-seq cells assigned to patch-enriched and matrix-enriched supertypes. As in **a**, cells mapping to T-type 0979 STR D2 Gaba 4 have been separated from the 0277 STR D2 Gaba 4 column and shown in their own column (0979). **c**, MERFISH section; each MSN is colored by patch/matrix score, predicted from regression to log(*Kremen1*) + log(*Sema5b*) - log(*Id4*) (see Methods). **d**, Kernel density estimates (KDEs) of patch/matrix score in Patch-seq cells assigned to supertypes (and T-type 0979) described in **a**. **e**, Scatterplot and fitted regression showing modest correlation between predicted patch/matrix score and input resistance in Patch-seq cells (top). Barplots showing input resistance values for matrix-enriched or patch-enriched supertypes (and T-type 0979) described in **a** (bottom). **f**, UMAP based on principal components of gene expression of 79,126 dissociated and 601 Patch-seq cells in STR D1 Gaba (D1 MSN), STR D2 Gaba (D2 MSN) and STR D1 Sema5a Gaba (D1 Sema5a MSN) subclasses, highlighting peripheral location of 0279 STR D2 Gaba 6 supertype, colored by T-type. **g**, Gene expression profile of Patch-seq cells assigned to T-types in 0279 STR D2 Gaba 6 supertype, normalized to maximum expression of each gene across all D2 MSNs. All other D2 MSNs are aggregated in rightmost column. **h**, MERFISH section containing CP and ACB, with soma locations of 0279 STR D2 Gaba 6 cells, colored by T-type. **i**, Input resistance of 0279 STR D2 Gaba 6 Patch-seq cells by T-type, compared to other D2 MSNs (gray). **j**, Subthreshold voltage responses of 0279 STR D2 Gaba 6 Patch-seq cells, averaged by T-type and compared to other D2 MSNs (gray). **k**, Rheobase of 0279 STR D2 Gaba 6 Patch-seq cells assigned to STR D2 Gaba 6 T-types, compared to other D2 MSNs (gray). **l**, Average relationship between firing frequency and stimulus intensity (f-I curves). **m**, UMAP in **f**, highlighting D1 Sema5a MSNs, colored by supertype. **n**, Gene expression profile of Patch-seq cells assigned to supertypes in D1 Sema5a subclass, normalized to maximum expression of each gene across MSNs. **o**, Same as **h**, containing D1 Sema5a MSN somata, colored by transcriptomic supertype. **p**, Comparison of input resistance for MSN subclasses. **q**, Averaged subthreshold voltage responses for MSN subclasses. **r**, Rheobase measurements for MSN subclasses. **s**, Average relationship between firing frequency and stimulus intensity (f-I curves). **t**, Example 0279 STR D2 Gaba 6 dendritic morphologies, colored by T-type. **u**, Example D1 Sema5a dendritic morphologies, colored by T-type.

We calculated a patch/matrix score for each neuron based on gene expression (Methods) for the Patch-seq dataset as well as the MERFISH data to confirm the association with compartmental structure (Fig. 3c). In Patch-seq data, the patch/matrix score separated representative patch-like and matrix-like MSN supertypes/T-type (Fig. 3d, Extended Data Fig. 7a). We found that input resistance and rheobase were modestly correlated with the patch/matrix score, suggesting higher input resistances and lower rheobases in patch-like MSNs^34^, which would be consistent with increased sensitivity to cortical inputs (Fig. 3e, top, Extended Data Fig. 7b; d, top). Input resistance varied significantly across patch-like and matrix-like MSNs (one-way ANOVA, F(4, 161) = 63.55, *p <* 0.001), with D1 patch-like MSNs exhibiting higher values than both D1 and D2 matrix-like MSNs, and neurons in the D2 patch-enriched supertype 0279 showing elevated input resistance compared to each other test group (all *p <* 0.001, Fig. 3e, bottom). Several dendritic morphological features, such as number of branches and total dendritic length, were correlated with patch/matrix score, with more dendritic branching apparent in matrix-like MSNs (Extended Data Fig. 7c; d, bottom). Interestingly, patch–matrix separation was not reflected as clearly as spatial location or D1/D2 identity in the LFs derived from MSN electrophysiological or morphological features; instead, the patch/matrix score was partially correlated with several LFs (Extended Data Fig. 7e).

As mentioned above, the patch-like supertype 0279 STR D2 Gaba 6 had distinctive properties in multiple modalities compared with other MSNs. The 0279 supertype appears at the periphery of the D2 region of the MSN transcriptomic UMAP and includes three T-types: 0986, 0987, and 0988 (Fig. 3f). Patch-seq cells from 0988 expressed *Htr7*, *Tac1*, and *Agtr1a*, corresponding to previously described *Htr7+* MSNs^3^ (Fig. 3g). 0986 cells expressed *Htr7*, *Tac1*, and *Th*, which, along with its ventral expression pattern in MERFISH data (Fig. 3h), makes it a strong candidate for being the T-type corresponding to the ”D2H MSNs” described in the ventromedial patch of the ACB medial shell^4^. 0987 expressed lower levels of *Htr7* and *Tac1* and had a patch-like spatial pattern (Fig. 3g-h)^27^. We asked whether these non-canonical MSNs also had unusual physiological properties and found that 0279 STR D2 Gaba 6 MSNs exhibited significantly higher input resistance (Welch’s t-test, *t* = 9.24, *p* = 2.8*e* − 10), lower rheobase (*t* = −12.16, *p* = 5.7*e* − 22) and consequent leftward shift in their firing rate–current (f–I) curves relative to other neurons in the D2 subclass (Fig. 3i-l). Together, these features suggest that 0279 STR D2 Gaba 6 neurons may require less synaptic input to evoke AP firing compared with other D2 MSNs.

D1 Sema5a MSNs are another unusual population of MSNs that are transcriptomically distinct from canonical D1 or D2 MSNs^27^(Fig. 3m) and co-express *Drd1* and *Drd2* and additional marker genes such as *Otof*, *Tshz1*, *Casz1*, and *Adarb2* (Fig. 3n). In MERFISH data, D1 Sema5a MSNs can be found throughout striatum, although supertype 0281 was enriched in CP, whereas the remaining supertypes were enriched in ACB (Fig. 3o). Cells mapping to this subclass also demonstrated a more excitable electrophysiological phenotype compared to canonical D1 and D2 MSNs (Fig. 3p-s), with higher input resistances and lower rheobases. Morphological reconstructions of D1 Sema5a MSNs suggest that they may have simpler and smaller dendritic arbors than D1 and D2 MSNs (Fig. 3u), which would be consistent with electrophysiological differences like higher input resistances; however, these observations are only preliminary given the small number of morphologies reconstructed for this subclass (*n* = 3).

### Cross-species conservation of continuous MSN gradients

We next asked whether the continuous cross-modal organization characterized in mouse MSNs was conserved in primates, given the importance for understanding striatal function in a translational context. Recent cross-species comparisons of striatal cell types describe overall similarity between mouse and non-human primate MSNs but also significant differences in marker gene expression and physiological properties like spike rate adaptation, latency to spike, membrane time constant and spike shape^35–37^. To investigate directly whether the variations and cross-modal correspondences in mouse MSNs are relevant in primates, we compared these findings with macaque MSNs from a Patch-seq dataset collected using very similar protocols^37^. We used the HMBA Consensus Macaque Basal Ganglia taxonomy^38^ to assign homologous macaque “Group” levels of the taxonomy to mouse types (Fig. 4a). A UMAP co-embedding of the combined mouse and macaque Patch-seq transcriptomic data showed that the two datasets were distinguishable in transcriptomic space but retained similar relationships between homologous transcriptomic groups (Fig. 4b).

**Figure 4:**
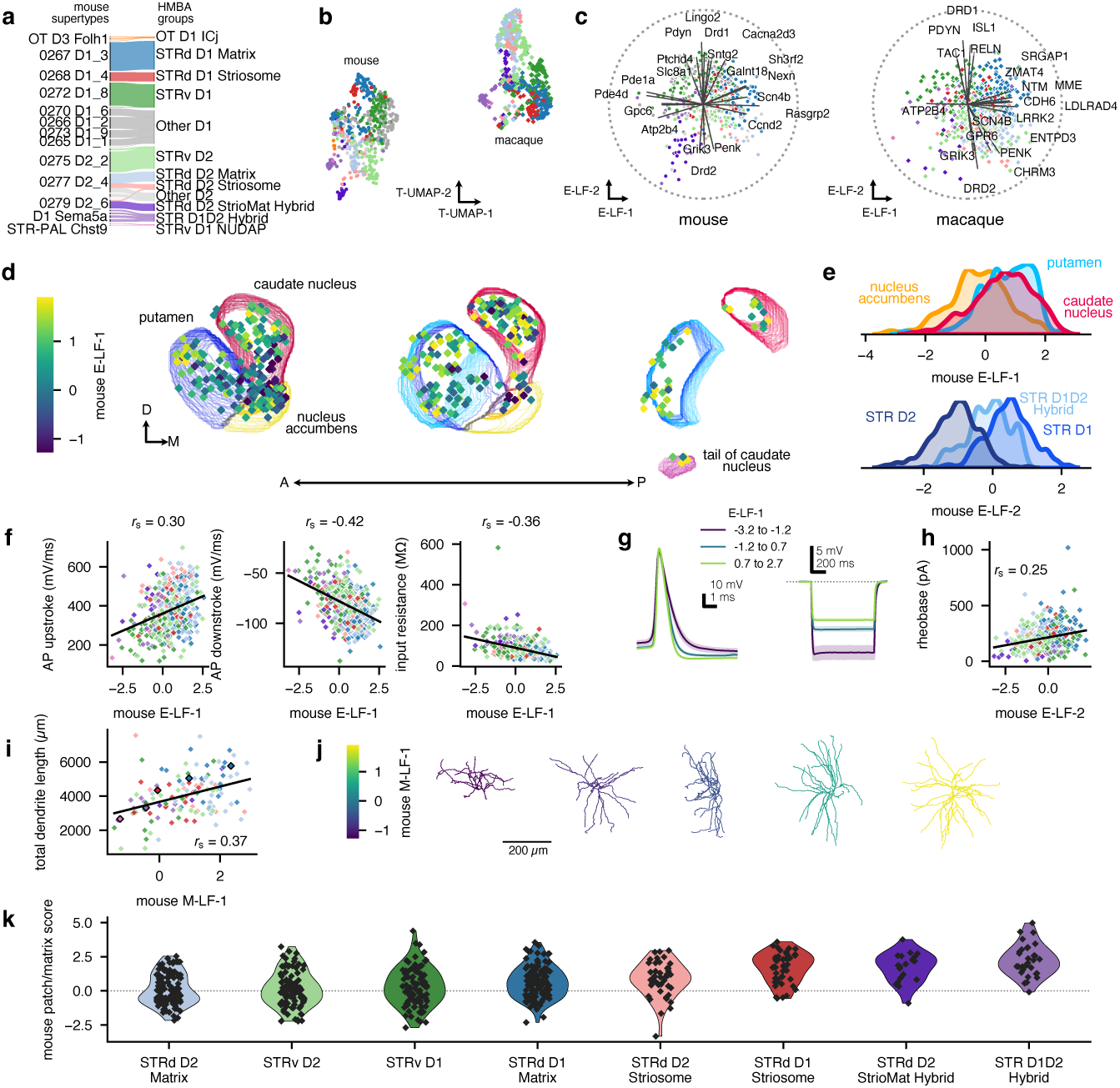
Cross-species comparison of cross-modal correspondences in MSNs. **a**, River plot showing HMBA Basal Ganglia Consensus taxonomy group assignments of homologous transcriptomic groups for mouse Patch-seq MSNs. ”STR” and ”Gaba” have been removed from mouse labels for brevity. **b**, UMAP embedding of gene expression from combined mouse (*n* = 624, circles) and macaque (*n* = 550, diamonds) MSNs reveals separation by species but also similarities in the relationships between homologous transcriptomic groups. **c**, Bi-plots of the first two electrophysiology latent factors (calculated from gene expression values). Mouse (left) and macaque (right) genes with the highest latent factor correlations are shown, and cells are colored by HMBA group. A correlation value of one is indicated by the dotted circle. **d**, Locations of Patch-seq cells in the average macaque template in three successive rostral-to-caudal sections. Somata from macaque cells (*N* = 478) are colored by their mouse E-LF-1 value. **e**, Kernel density estimates (KDEs) of predicted mouse E-LF-1 (top, colored by structure) and E-LF-2 (bottom, colored by HMBA subclass) distributions for macaque Patch-seq cells. **f**, Relationship between mouse E-LF-1 and AP upstroke (left) downstroke (middle) and input resistance (right) for macaque cells. Lines show a linear fit, and Spearman correlations are reported. **g**, Averaged action potential waveforms (left) and averaged subthreshold voltage responses (right) for macaque cells grouped by mouse E-LF-1. **h**, Relationship between mouse E-LF-2 and rheobase for macaque cells. The line represents a linear fit to the data. **i**, Relationship between mouse M-LF-1 and total dendritic length for macaque dendritic reconstructions. The line represents a linear fit to the data. Outlined markers indicate the examples shown in (**j**). **j**, Examples of dendritic macaque MSN morphologies, ordered and colored by mouse M-LF-1 value. **k**, Distributions of predicted patch/matrix scores in macaque MSNs, across HMBA groups. Cells assigned to striosome and matrix associated HMBA groups have scores on opposite ends of the distribution.

To investigate whether the same phenotypically linked transcriptomic gradients are present in the macaque dataset, we applied weights from mouse sparse RRR to homologous genes in macaque (491 homologous genes out of 557 genes with non-zero weights for E-LFs) to estimate mouse-derived latent factors for the macaque cells. Mouse and macaque HMBA groups occupied similar locations in the latent factor space defined by the mouse E-LF-1 and E-LF-2, with some similarities in the most correlated genes, in particular direct/indirect pathway associated genes *Drd1*, *Drd2*, *Pdyn* and *Penk*, as well as sodium channel auxiliary subunit gene *Scn4b* and the calcium pump gene *Atp2b4* (Fig. 4c). Macaque neurons located in the ACB had lower E-LF-1 values, paralleling the spatial variation seen in mouse (Fig. 4d-e). Macaque MSNs also had similar relationships between these latent factors and electrophysiological features: macaque cells with higher mouse E-LF-1 values exhibited faster AP kinetics and had lower input resistances (Fig. 4f-g). The mouse E-LF-2 separated macaque cells from the D1 and D2 HMBA subclasses and was associated with variation in neuronal excitability (Fig. 4e, h). These parallels extended to morphological features as well (calculated with 136 homologous genes out of 149 genes with non-zero weights for M-LFs): macaque neurons with higher mouse M-LF-1 values had larger dendritic arbors (Fig. 4i-j). Finally, we applied the mouse-derived patch/matrix score regression model to homologous genes in macaque (1,036 homologous genes out of 1,210 original mouse genes) and found that striosome-identified macaque subclasses were at one end, and matrix-identified subclasses were at the other (Fig. 4k); the StrioMat Hybrid and D1D2 Hybrid groups also had high striosome-like patch/matrix scores. Together these findings suggest that, despite evidence of divergence in a number of physiological properties and gene expression between mouse and macaque MSNs, broad gene programs and major features of MSN organization appear to be conserved across these two species.

### Multimodal characterization of striatal interneurons

Next, we used our Patch-seq dataset of mouse striatal interneurons to test whether coordinated variation between transcriptomic identity and physiological properties generalizes beyond MSNs. Although interneurons represent only ∼5% of cells in the mouse striatum, they display considerable diversity^7,39,40^. Our Patch-seq dataset includes cells mapping to the transcriptomic supertypes 0234 STR Prox1 Lhx6 Gaba 2 (containing the T-types 0833, 0834, and 0836), 0260 PAL-STR Gaba-Chol 2 (containing the T-types 0929, 0930, 0931, and 0933), and multiple T-types within the STR Lhx8 Gaba (including 0840 and 0842), Sst Chodl Gaba (including 0849) and Vip Gaba (including 0639 and 0641) subclasses (Fig. 5a). Projecting dissociated and Patch-seq neurons into a UMAP space showed that striatal interneuron supertypes mostly occupy distinct locations (Fig. 5b). These transcriptomic groups differentially expressed key marker genes for a variety of striatal interneuron types (Extended Fig. 8a, right). Overall, the morpho-electric properties of the neurons were consistent with previous descriptions of transcriptomically-defined cell types (Fig. 5c). For example, 0234 STR Prox1 Lhx6 Gaba 2 and 0260 PAL-STR Gaba-Chol 2 aligned well with previous descriptions of fast-spiking and cholinergic interneurons, respectively^39,41^. Using cells for which we had triple-modality data, we also confirmed that these different sets of interneurons were grouped into distinct MET-types that had one-to-one or several-to-one relationships with T-types (Extended Data Fig. 1).

**Figure 5:**
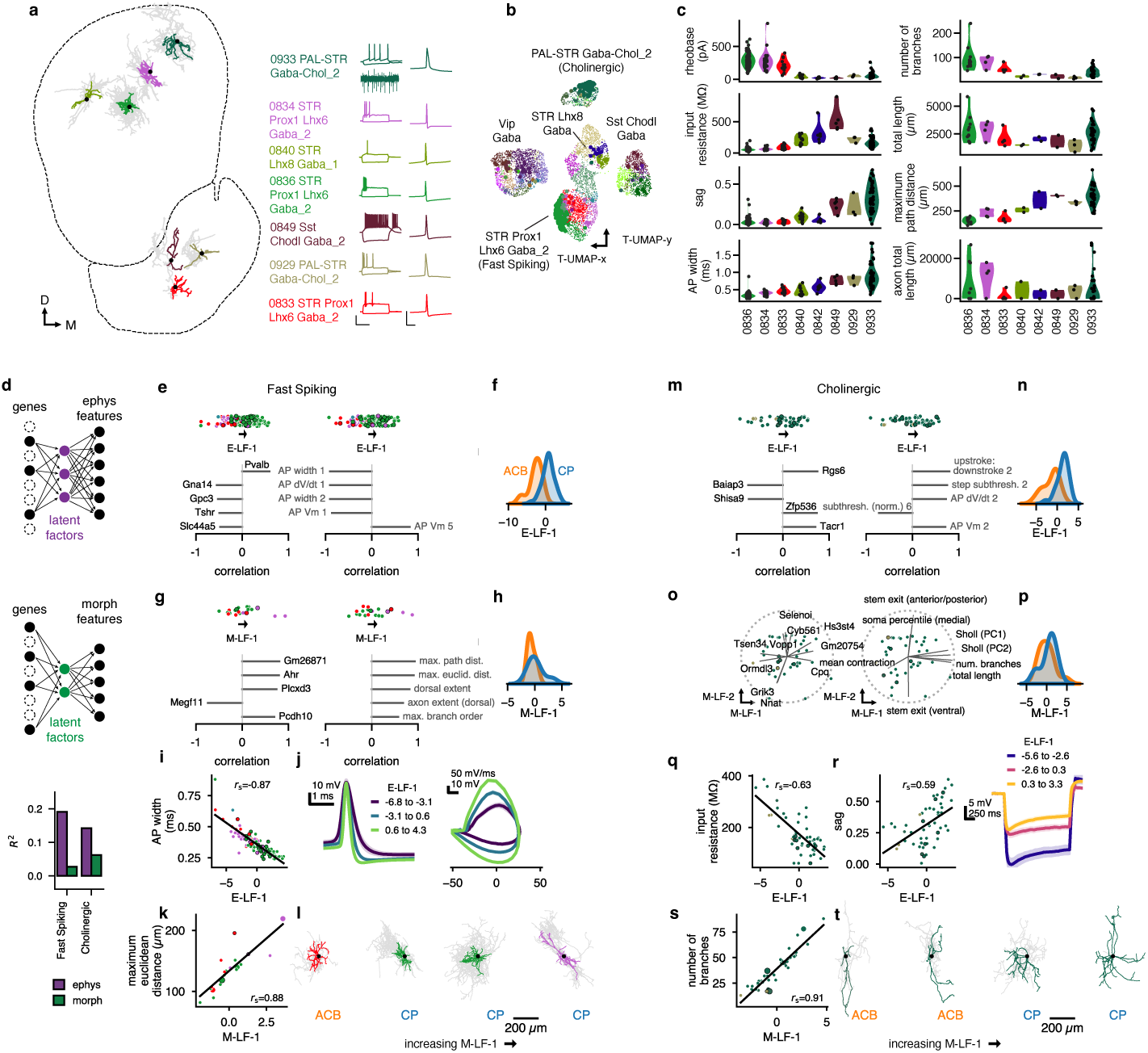
Discrete and continuous variation in striatal interneuron properties. **a**, Coronal section of CP and ACB in Allen CCFv3 with example registered Patch-seq reconstructions (dendrites are colored by T-type, axon is in gray) and current clamp recordings (left: subthreshold and suprathreshold responses to long (1 s) current step, right: single action potential (AP) from short (3 ms) current step). A cell-attached recording (1 s) is shown for the example cholinergic interneuron, demonstrating spontaneous firing. **b**, UMAP of gene expression of reference dissociated cells (small markers, n=6,225) and Patch-seq cells (large markers, n=395), colored by T-type. **c**, Electrophysiological (left) and morphological (right) characteristics of striatal interneuron T-types. **d**, Schematics of RRR for electrophysiology (ephys; top) and morphology (morph; middle) features, and cross-validated *R*^2^ values for prediction of ephys (purple) and morph (green) (top). RRR was run separately for the two modalities and for Fast Spiking and Cholinergic interneurons. **e**, Genes (left) and features (right) with the highest correlations to the sole ephys latent factor (LF) in Fast Spiking interneurons. Color indicates T-type; black borders indicate held-out cells not used to fit the regressions. **f**, Kernel density estimates (KDEs) of E-LF-1 distributions in ACB and CP Fast Spiking interneurons. **g**, Genes (left) and features (right) with the highest correlations to M-LF-1 in Fast Spiking interneurons. **h**, KDEs of M-LF-1 distributions in ACB and CP Fast Spiking interneurons. **i**, Scatterplot and fitted regression showing correlation of E-LF-1 and AP width in Fast Spiking interneurons. **j**, Averaged AP waveforms and AP phase plots for Fast Spiking interneurons grouped by E-LF-1 value. **k**, Scatterplot and fitted regression showing correlation of M-LF-1 to dendrite maximum euclidean distance in FS cells. Larger discs denote cells in **m**. **l**, Example morphological reconstructions of FS cells, spanning the range of M-LF-1 values. Dendrites are colored by T-type, local axon is in gray. **m**, Genes (left) and features (right) with the highest correlations to E-LF-1 in Cholinergic cells. Color indicates T-type; black borders indicate held-out cells not used to fit the regressions. **n**, KDEs of E-LF-1 distributions in ACB and CP FS cells. **o**, Bi-plot of M-LF-1 and M-LF-2 for Cholinergic cells, with most highly correlated genes (left) and features (right). **p**, KDEs of M-LF-1 distributions in ACB and CP Cholinergic cells. **q**, Scatterplots and fitted regressions showing correlations of E-LF-1 with input resistance (left) and hyperpolarization-induced sag ratio (right) in ChAT cells. **r**, Averaged subthreshold voltage responses for Cholinergic cells grouped by E-LF-1 value. **s**, Scatterplot and fitted regression showing correlation of M-LF-1 with number of dendritic branches in Cholinergic cells. Larger discs denote cells in (**u**). **t**, Example ChAT cell morphological reconstructions, spanning the range of M-LF-1 values. Dendrites are colored by transcriptomic type, local axon is in gray.

Fast-spiking striatal interneurons mapped to multiple *Pthlh+* T-types (0833, 0834, 0836) in 0234 STR Prox1 Lhx6 Gaba 2 supertype and as a group had the narrowest AP widths of striatal interneurons (0.36 ms ± 0.08 ms, *n* = 148; Fig. 5c, left). Together, these T-types formed the STR Prox1 Lxh6 MET-type (Extended Data Fig. 1b, c). They had basket cell-like morphologies with compact dendritic fields (Fig. 5a). 0836 locations were strongly biased toward CP and had higher expression of *Pvalb* than 0833, which was primarily found in ACB; cells mapping to 0834 were found in both ACB and CP (Extended Data Fig. 8a,b).

Cholinergic interneurons exhibited wider APs than most other interneurons (0.90 ms ± 0.33 ms, *n* = 53), varying levels of hyperpolarization-induced sag, spontaneous firing at rest, and large morphologies with extensive axonal fields (Fig. 5a,c). Most of the striatal cholinergic interneurons in our dataset mapped to a single T-type (0933), with a handful of cells mapping to other related T-types (0929, 0930, and 0931) that appeared more often in ventral striatum (Extended Data Fig. 8a). These cells also were grouped into a single MET-type (STR Chol, Extended Data Fig. 1b, c).

We also collected examples of other striatal interneurons, albeit with less extensive sampling. The transcriptomic subclass STR Lhx8 Gaba likely corresponds to non-dopaminergic, GABAergic *Th+* expressing neurons (THINs)^39,40,42^ as they are the only striatal interneuron population expressing *Th* (Fig. 5a, Extended Data Fig. 8a, b). These cells mapped to two primary T-types (0840 and 0842 STR Lhx8 Gaba 1) along with a few other related T-types and formed a single MET-type (STR Lhx8, Extended Data Fig. 1b, c). This group had an average AP width intermediate between fast-spiking and cholinergic interneurons (0.49 ms ± 0.14 ms, *n* = 25), and demonstrated some variation in input resistance and AP dynamics across *Th+* T-types (Fig. 5c). The 0239 Sst Chodl Gaba 2 supertype likely corresponds to *Sst*+ low-threshold spiking (LTS) interneurons. We identified a small number of these *Sst+ / Chodl+ / Nos1+ / Npy+* cells in our Patch-seq data (Extended Data Fig. 8a, d), with the highest input resistances of striatal interneurons (497.88 MΩ ± 203.48 MΩ, *n* = 9) moderate amounts of hyperpolarization-induced sag, and a hallmark Ca2+ dependent low-threshold spike in response to depolarization, or rebound firing after a hyperpolarizing current injection (Fig. 5a,b). We also found 38 cells that mapped to the Vip Gaba subclass (Extended Data Fig. 8a, e). While T-types 0641 and 0639 were overrepresented, cells diffusely mapped to many T-types within the subclass. A subset of these co-expressed *Cck* and may be correlates of the rare, less distinct aspiny *Cck*+ neurogliaform cells reported in previous studies^7,43,44^. We then investigated how multimodal properties co-varied within the fast-spiking/fast-spiking-like and cholinergic interneuron populations, as these were the interneuron subclasses best represented in our dataset. As with MSNs (see Figure 2), we applied sparse RRR to their electrophysiology and morphology features separately, obtaining relatively higher performance for predicting electrophysiology features (Fig. 5d). Only one E-LF and M-LF was fit for fast-spiking interneurons. E-LF-1 was strongly linked to spatial location, AP kinetics, and expression of *Pvalb* and *Gna14*, a Gq-family G*_α_* subunit (Fig. 5e, f, i, j). Fast-spiking cells with low E-LF-1 values were more frequently found in ACB, mapped to 0833, and had slower AP upstroke and downstroke. This matched the relationship we characterized with E-LF-1 in MSNs (i.e., faster APs at the dorsolateral end of the gradient), so there appeared to be co-variation of these electrophysiological properties in the striatal microcircuits. The three major fast-spiking T-types (0833, 0834, 0836) also differed from each other along this main dimension (Fig. 5e). Fast-spiking M-LF-1 was associated with the maximum path length of dendritic processes and did not differentiate ventral from dorsal striatum (Fig. 5g, h, k, l).

Cholinergic interneurons also demonstrated robust transcriptomically linked variation in their properties. A single E-LF was fit from sparse RRR and was associated with subthreshold electrophysiological properties, including input resistance and hyperpolarization-induced sag, as well as AP shape and location within striatum (Fig. 5m, n, q, r). The top contributing genes to E-LF-1 included *Rgs1* (a regulator of G protein signaling), *Shisa9* (an auxiliary AMPA receptor subunit). There was a strong association between M-LF-1 and the complexity of dendritic arbors in cholinergic interneurons (Fig. 5s,t).

### Multimodal characterization of cell types in GPe, STN, GPi, SNr, and SNc/VTA

Next, we used our Patch-seq dataset to characterize and validate cell types in basal ganglia structures other than the striatum (Fig. 6a-e). In GPe, cells in MGE-derived supertype 0247 NDB-SI-MA-STRv Lhx8 Gaba 4 corresponded to prototypic neurons^27^, which are known to project to other basal ganglia structures and express *Nkx2.1*, *Pvalb* and *Lhx6*^27,45–47^ (Fig. 6f). Cells in the GPe from this subclass most frequently mapped to 0892 but also to several other related T-types. Prototypic cells were spontaneously active at rest and had narrow APs (0.38 ms ± 0.12 ms, *n* = 16), and could sustain high firing frequencies (Fig. 6a, g)^47,48^. They also exhibited high membrane capacitance (103.37 pF ± 27.99 pF), consistent with their large soma surface area (837.35 µm^2^ ± 595.02 µm^2^, *n* = 4)^48,49^. We found that cells mapping to the LGE-derived supertype 0257 GPe-SI Sox6 Cyp26b1 Gaba 1^27^ expressed *Foxp2*, *Npas1*, *Meis2*, and *Penk* and corresponded to arkypallidal neurons^27^, which project to striatum^45–47,49^ (Fig. 6a, f). Arkypallidal neurons had slightly broader (0.47 ms ± 0.10 ms, *n* = 7) APs and lower effective membrane capacitance (66.08 pF ± 17.28 pF) than prototypic neurons (Fig. 6a, g)^48^. Morphologies reconstructed from arkypallidal recordings had smaller cell bodies (438.37 µm^2^ ± 15.01 µm^2^, *n* = 2) and dendritic arbors than prototypic cells^48^ (Fig. 6h). Finally, we encountered an example of a cell mapping to PAL-STR Gaba-Chol, which corresponded to a rarer GPe cell type — frontal cortex-projecting cholinergic cells that co-release GABA and acetylcholine^50,51^. This cell exhibited slower AP kinetics compared to the other GPe types (Fig. 6a (iv)).

**Figure 6:**
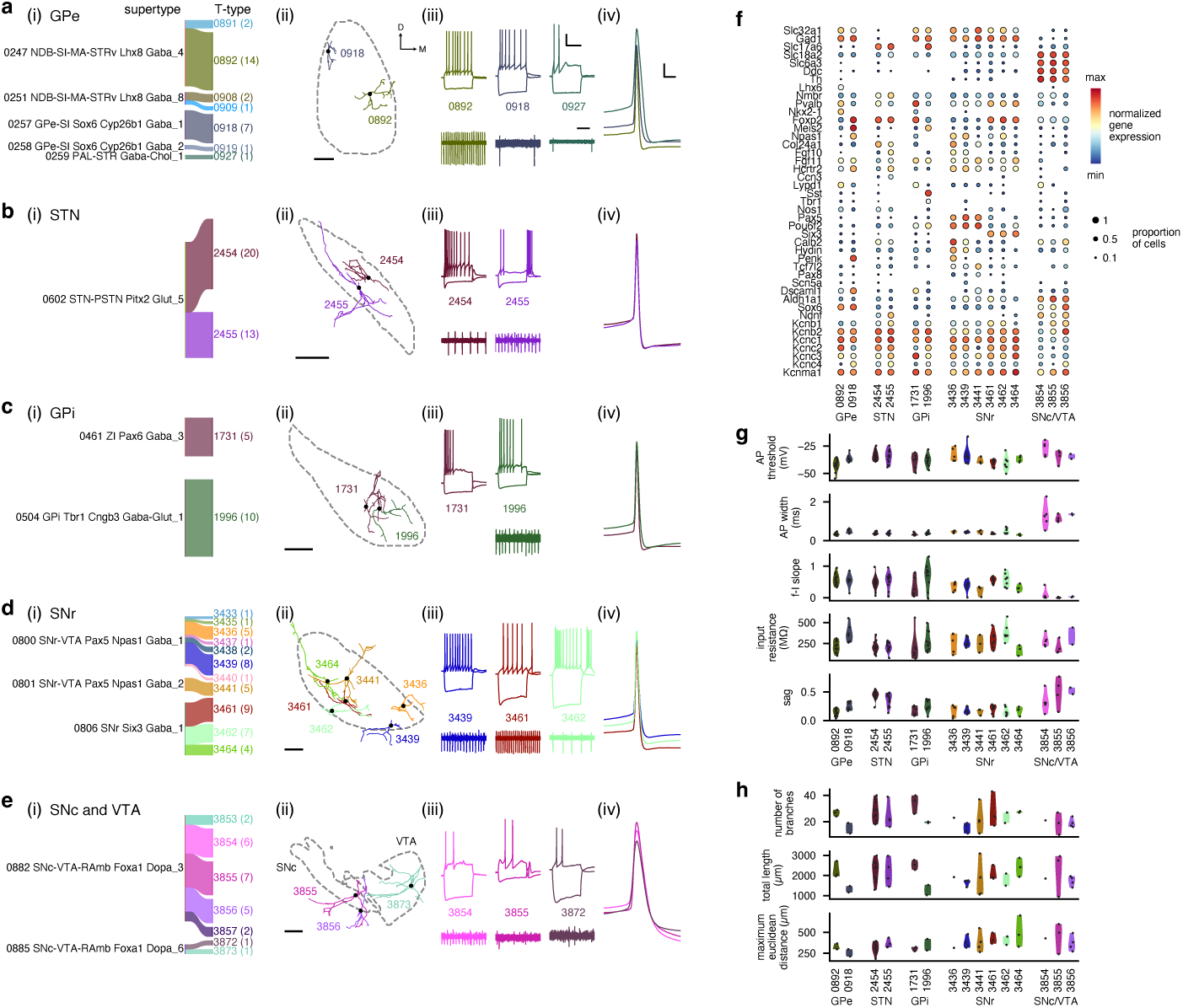
Multimodal characterization of cell types in GPe, STN, GPi, SNr, and SNc. **a**, (i) Supertypes (left) and T-types (right) of cells in the GPe. Numbers in parentheses indicate number of cells that have transcriptomic data as well as electrophysiology and/or morphology data. (ii) Example morphologies of cells in GPe (structure outlined with dotted line). Scale bar: 200 µm. (iii) Example electrophysiological responses to long current steps (top) and example cell-attached recordings showing spontaneous firing (bottom) for cells in different T-types. Colors match sub-panel (i). Top scale bar: 20 mV, 500 ms. Bottom scale bar: 100 ms. (iv) Example action potential waveforms. Colors match sub-panel (i). Scale bar: 10 mv, 1 ms. **b**, Same as (**a**) but for cells in STN. **c**, Same as (**a**) but for cells in GPi. Note that a cell-attached trace was not available for the example shown for T-type 1731. **d**, Same as (**a**) but for cells in SNr. **e**, Same as (**a**) but for cells in SNc and VTA. **f**, Gene expression patterns of cell types in basal ganglia structures downstream of the striatum. Color indicates average expression, normalized to the maximum expression of that gene among all included cells. Size of dot indicates the fraction of cells with non-zero expression of that gene. **g**, Electrophysiological characteristics of cell types in basal ganglia structures downstream of the striatum. **h**, Morphological characteristics of cell types in basal ganglia structures downstream of the striatum.

STN Patch-seq cells in our dataset mapped to 2 closely related glutamatergic T-types: 2454 and 2455 STN-PSTN Pitx2 Glut 5 (Fig. 6b). The STN T-types did not show robust differences in *Pvalb* or *Col24a1* expression in the Patch-seq data (Fig. 6f), but 2454 revealed a modest but significant increase in hyperpolarization-induced sag compared to 2455 (Welch’s t-test, *t* = 2.36, *p* = 0.028), which overall is consistent with previous reports^52^ (Fig. 6g). Finally, STN morphologies were a largely homogeneous group, with a slight trend towards longer dendrites (max path distance) in 2455 (Fig. 6h).

In the GPi, we identified the T-types for *Pvalb+* motor thalamus-projecting cells (1731 ZI Pax6 Gaba 3) and *Sst+* lateral habenula (LH)-projecting cells (1996 GPi Tbr1 Cngb3 Gaba-Glut 1)^8^. *Pvalb+* Patch-seq cells also co-expressed *Lypd1*, another marker of thalamus-projecting cells^8^, and had fast APs (0.32 ms ± 0.08 ms, *n* = 5). *Sst+* Patch-seq cells co-expressed the expected markers *Slc17a6* and *Tbr1*^8^, had slightly broader APs (0.38 ms ± 0.07 ms, *n* = 10), and also exhibited hyperpolarization-induced sag (0.26 ± 0.10 vs 0.17 ± 0.08, NS). Despite the limited number of reconstructed cells, preliminary observations suggested that *Pvalb+* neurons (1210.32 µm^2^ ± 316.39 µm^2^ surface area, 2530.82 µm ± 245.95 µm dendritic length, *n* = 3) had larger somata and more extensive dendritic arborizations than *Sst+* neurons (738.92 µm^2^ ± 62.33 µm^2^ surface area, 1276.75 µm ± 360.22 µm dendritic length, *n* = 2) (Fig. 6h).

Our Patch-seq data from SNr mapped to several T-types within three supertypes (Fig. 6d). Based on marker gene expression and ventromedial localization in SNr, cells mapping to supertypes 0800 SNr-VTA Pax5 Npas1 Gaba 1 and 0801 SNr-VTA Pax5 Npas1 Gaba 2 correspond to previously described *Pou6f2+/ Pax5+* cells, whereas cells mapping to supertype 0806 SNr Six3 Gaba 1 correspond to previously described *Six3+ / Foxp2+* cells (Fig. 6f, d (ii))^23^. Both of these groups fired spontaneously and had narrow AP widths (0.38 ms ± 0.09 ms, *n* = 18 cells from 0806 SNr Six3 Gaba 1; 0.44 ms ± 0.08 ms, *n* = 22 cells from 0800 SNr-VTA Pax5 Npas1 Gaba 1 and 0801 SNr-VTA Pax5 Npas1 Gaba 2). Both groups had dendritic arbors that spanned large distances across the SNr (Fig. 6d (ii), h). We did not sample any cells mapping to a third reported *Tcf7l2+ / Scn5a+* SNr subclass^23^.

We also collected data from a small number of dopaminergic cells primarily in the SNc (*n* = 10) but also the VTA (*n* = 3), which mostly mapped to supertype 0882 SNc-VTA-RAmb Foxa1 Dopa 3 with a smaller number mapping to supertype 0885 SNc-VTA-RAmb Foxa1 Dopa 6. The cells in supertype 0882 SNc-VTA-RAmb Foxa1 Dopa 3 were *Sox6+*, *Alhd1a+*, and *Ndn4+*, linking them to a subset of dopaminergic neurons with increased vulnerability in a model of Parkinson’s disease^9^. The physiological characteristics of dopaminergic neurons were distinct from other sampled non-striatal types, with profoundly longer duration APs, large, clear voltage sag in response to hyperpolarization, and subthreshold oscillations (Fig. 6e, g).

Several morphoelectric properties, such as those relating to AP shape and dendritic morphology, were similar across types of neurons in these different basal ganglia structures, and we found that our MET-type classification combined neurons from the SNr, GPe, and GPi into two related MET-types (Extended Data Fig. 1b, c). There were some differences in the composition of the two MET-types, however: the SNr-GPe-GPe-1 MET-type had most of the cells from the SNr-VTA Pax5 Npas1 subclass along with the GPe-SI Sox6 Cyp26b1 Gaba 1 cells (arkypallidal GPe cells), while the SNr-GPe-GPe-2 MET-type had more cells from the SNr Six3 Gaba subclass, the NDB-SI-MA-STRv Lhx8 Gaba 4 supertype (prototypic GPe cells), and the two major GPi types. Glutamatergic cells from the STN and dopaminergic cells from the SNc and VTA each formed their own distinct MET-types (STN and SNc-VTA Dopa, respectively; Extended Data Fig. 1b, c).

Neurons in these structures (apart from SNc/VTA) generally had narrow APs with a rapid repolarization phase after the peak (Fig. 6a-e (iv)); consistent with this, we found high levels of expression of members of the *Kcnc* family (Fig. 6f), which encode Kv3 potassium channels associated with narrow action potentials and rapid firing. The particular *Kcnc* genes expressed did, though, vary by T-type. Also, neurons in the SNc/VTA that had much wider APs did not have high levels of *Kcnc* expression. Other potassium channel genes, such as *Kcnb1* and *Kcnb2* (Kv2 channels) and *Kcnma1* (BK channels) were also widely expressed; these channels have previously been shown to be involved in modulation of AP shape and firing patterns in SNc dopaminergic neurons^53^.

### MSN projection patterns to other BG structures

Given the systematic differences found among MSNs, we asked whether these organizing principles were preserved as information propagates through the basal ganglia circuit. Do MSN long-range projections maintain this primary dimension of transcriptomic and spatial variation, and are these channels of information kept separate in downstream targets, or do they get pooled together? Even if the axonal projections of MSNs are organized consistent with this major transcriptomic gradient, the dendrites of neurons in those target structures can span relatively large distances (see Figure 6a-e (ii)), which might allow them to pool inputs regardless.

To investigate how co-variation across transcriptomic identity, spatial location, and phenotype might be linked across different stages of the basal ganglia circuit, we analyzed 264 complete whole-neuron morphologies of CP MSNs from Peng *et al.* [54] that were reconstructed from whole brain fluorescence Micro-optical Sectioning Tomography (fMOST) imaging and registered to the Allen CCFv3. We determined whether each neuron was located within the recently-delineated lateral (CPl), dorsomedial (CPdm), ventromedial (CPvm), or intermediate-ventral (CPiv) CP subdivisions^22^ (Fig. 7a) — these CP subdivisions are associated with different sets of cortical and thalamic inputs. We also used the sparse RRR fits from the Patch-seq dataset to estimate morphological latent factor values for each neuron using the shared set of dendritic features between the two datasets (Fig. 7b-c). We then analyzed the MSN axon locations within their target structures (SNr, GPi, and GPe) and calculated both the distribution of axons arising from different subdivisions (Fig. 7d, f, h (ii)) and the average predicted M-LF-1 values of axons at different locations throughout the target structure (Fig. 7d, f, h (iii)).

**Figure 7:**
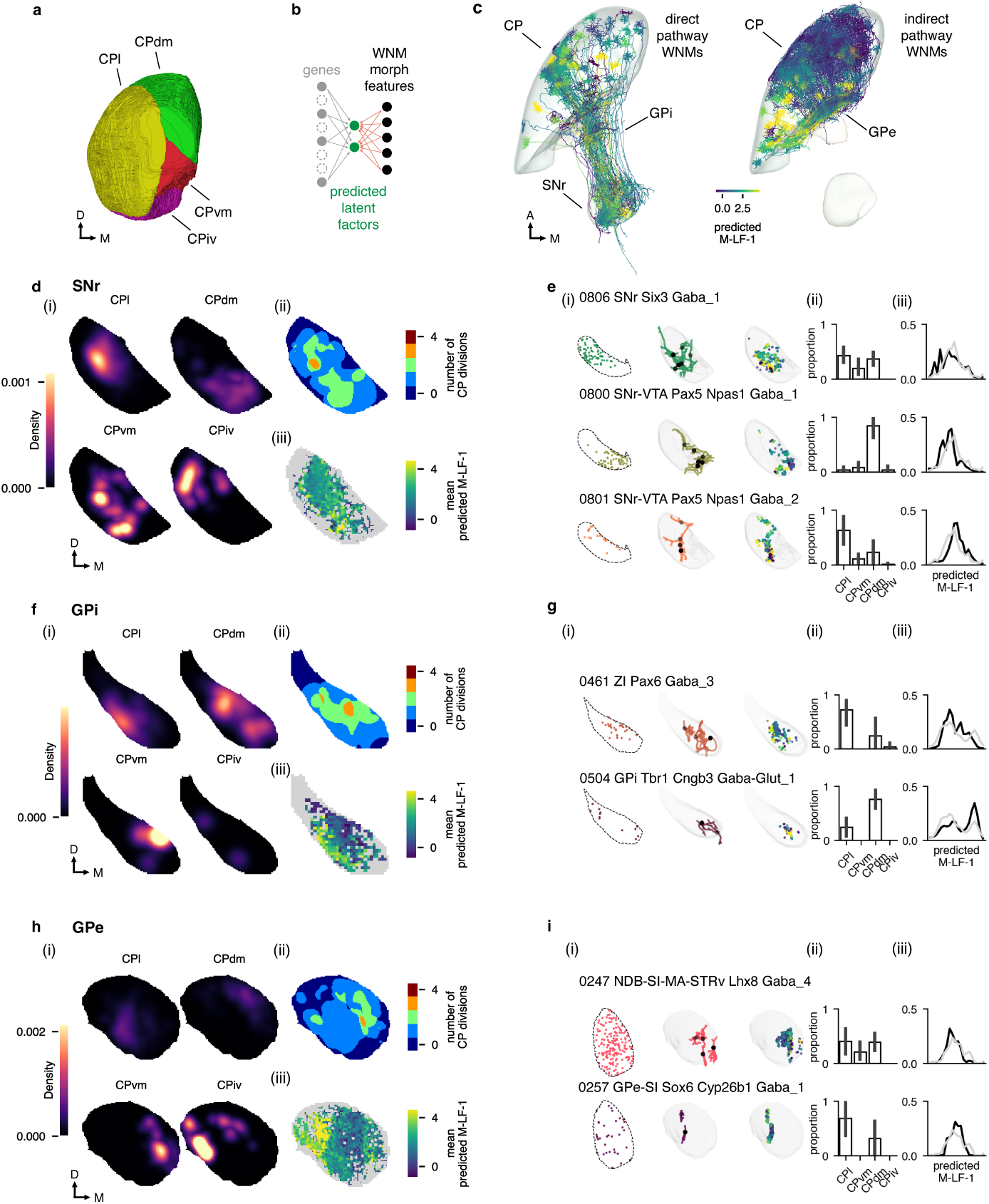
Relationship between MSN gradient and projection patterns. **a**, CP subdivisions in coronal view: lateral (CPl), dorsomedial (CPdm), ventromedial (CPvm), intermediate-ventral (CPiv) and posterior (CPp, not shown). **b**, Schematic showing prediction of morphological latent factors from whole neuron morphology (WNM) dendritic features using sparse reduced-rank regression fit on Patch-seq data. **c**, WNMs of cells projecting from CP to SNr and GPi (left) and projecting from CP to GPe (right). Cells are colored by predicted M-LF-1. **d**, (i) Coronal maximum projection density maps of axon terminals and branches in SNr arising from WMNs in CPl, CPdm, CPvm and CPiv. (ii) Mean predicted M-LF-1 in axon terminals and branches of WNMs in SNr. (iii) Diversity of CP subdivisions reflected in WNM axon terminals and branches in SNr. **e** (i) Dendritic sampling by transcriptomic supertypes in SNr. From left: MERFISH section showing spatial distribution of transcriptomic supertype; dendritic morphologies of Patch-seq cells mapping to supertype; WNM axon branches and nodes within 50 µm radius of Patch-seq dendrites. (ii) Relative proportions CP subdivisions represented by nearby WNM axons (iii). KDE of predicted M-LF-1 distribution in nearby WNM axons. **f**, Same as **d** but for GPi. **g**, Same as **e** but for GPi. **h**, Same as **d** but for GPe. **i**, Same as **e** but for GPe.

Neurons from different CP subdivisions projected to different locations within SNr (Fig. 7d), GPi (Fig. 7f), and GPe (Fig. 7h). For example, MSNs in CPl projected to dorsolateral and central SNr, while MSNs in CPdm projected to ventromedial SNr. These inputs were relatively segregated; most locations within these structures were primarily innervated by only one or two CP subdivisions (Fig. 7d, f, h - part ii).

Interestingly, the M-LF-1 gradient was preserved in axonal projection patterns to different extents in different structures. The MSN axons in SNr did not appear to be strongly organized by predicted M-LF-1 (Fig. 7d-iii). On the other hand, axons in GPi and GPe exhibited larger variation in predicted M-LF-1 across the structure, with higher values closer to the edges of GPi (Fig. 7f-iii) and at the more medial aspect of GPe (Fig. 7h-iii).

Next, we were interested in whether there were cell-type specific differences in the areas that downstream neurons might sample from. We used the reconstructed morphologies of Patch-seq neurons registered to the Allen CCFv3 and identified the MSN axons near those dendrites (Methods). We then analyzed the CP subdivision origins of those neurons as well as their predicted M-LF-1 values. In the SNr, different transcriptomic supertypes are located in different parts of the SNr as shown both by MERFISH data and the registered locations of Patch-seq neurons (Fig. 7e-i). Even though the dendrites of these neurons are elongated, we still found that there were supertype-specific differences in the CP subdivisions from which they could be receiving inputs (Fig. 7e-ii)—neurons in the 0800 supertype had dendrites positioned to receive inputs from CPdm axons, while the 0801 supertype had its dendrites nearer to CPl axons. In contrast, all three SNr supertypes examined showed little difference in the distribution of M-LF-1 values of the axons their dendrites were near (Fig. 7e-iii), which was unsurprising given that axons with different M-LF-1 values were not particularly segregated with the SNr (Fig. 7d-iii).

Cell types in the GPi, however, differed both in terms of the CP subdivision and M-LF-1 values of potential inputs. Neurons in the 0461 ZI Pax6 Gaba 3 supertype, found in the GPi core (Fig. 7g-i), had their dendrites nearer axons from the CPl subdivision than the CPdm subdivision (Fig. 7g-ii) and were nearer axons with lower M-LF-1 values on average (Fig. 7g-iii). Neurons in the 0504 GPi Tbr1 Cngb3 Gaba-Glut 1 supertype, found in the GPi shell, exhibited the opposite pattern (Fig. 7g). There were also differences in potential CP subdivision innervation among supertypes in the GPe (Fig. 7i (i-ii)), but the distributions of M-LF-1 values of nearby axons were fairly similar (Fig. 7i (iii)). We do note, though, that we did not have any morphological reconstructions at the lateral edge of GPe where projecting axons had the highest M-LF-1 values. Overall, the topographic organization of MSN projections appeared to differ across downstream structures, with the transcriptomic gradient preserved to the largest extent in GPi and the least in SNr.

## Discussion

Large-scale, high-throughput single cell RNA sequencing and spatial transcriptomics have reshaped our view of cell types in the basal ganglia, simultaneously confirming known distinctions across cell types while also uncovering new axes of genetic and spatial variation. However, it remains unclear how this observed molecular diversity translates into cellular phenotypes and, ultimately, circuit-level functions. Here we used the Patch-seq technique to characterize the electrophysiological and morphological properties of T-types at the individual cell level, providing a framework for linking traditional cell classifications to the transcriptomic taxonomy while also describing cross-modal variation within cell types. Our dataset spanned the striatum as well as multiple other BG structures, providing a fuller view of the variation within and across BG cell types; in addition, the transcriptomic data from recorded Patch-seq neurons allowed us to reference MERFISH spatial transcriptomics data by virtue of the shared transcriptomic taxonomy. The morphological features measured from Patch-seq neurons also allow us to connect these data with other morphological datasets, such as the whole-neuron morphology data we used to analyze MSN projection patterns.

### Variation across MSNs

Molecular genetics has expanded the landscape of MSN diversity and moved beyond the traditional direct/indirect pathway framework, but a key challenge remains in linking this molecular heterogeneity to phenotype and function. MSNs are genetically separable into dozens of T-types^2^, but interpreting the functional significance of this diversity requires multimodal approaches. Much of the molecular diversity in MSN T-types corresponds with their positions within the striatum^4,27^. Here we observe a similar pattern in our Patch-seq data, with MSN T-types tiling the continuous electrophysiological and morphological latent spaces defined by sparse RRR. Notably, the ventromedial-dorsolateral spatial axis emerged as the primary axis of transcriptomic, spatial, and phenotypic variation in MSNs, whereas D1 vs D2 (historically the most common basis for classifying these cells) appeared as an orthogonal axis. We also demonstrate that this organizing structure (encompassing gene expression, location, electrophysiology, and morphology) was also consistent between mouse and macaque MSN populations, suggesting that many fundamental features of the circuit are well-conserved even while certain specific morphoelectric features^37^ or marker genes^38^ may vary more dramatically between the two species.

Our work adds additional detail to the continuous spatio-molecular gradient in striatum for which there is already robust evidence^3,4,21^. Linking this gradient to continuous variation in the morphoelectric properties of MSNs suggests that they may be coordinated to jointly shape how MSNs along the ventromedial to dorsolateral axes integrate synaptic inputs and convey information to downstream basal ganglia structures. MSNs integrate multiple converging inputs from cortex, thalamus, local interneurons, in addition to neuromodulatory inputs^55^. Broadly, CP processes sensorimotor information, whereas ACB is more strongly involved in motivational aspects of learning^56^, and inputs are further organized in multiple parallel, topographical channels^18,19,22^. We encountered the fastest AP kinetics, and largest, most complex dendritic arbors in MSN T-types found toward the dorsolateral striatum. Precision of AP timing may be particularly important for shaping action selection in dorsal striatum, and expansive dendritic morphologies may enable MSNs to sample more widely, potentially integrating synaptic inputs from different functional domains. Conversely, limbic processing in the ventromedial striatum may not only be tolerant of slower AP dynamics, but in fact benefit from integrating contextual inputs over time. The multimodal continuum of MSNs appears mirrored in the AP kinetics of the nearby fast-spiking striatal interneurons. It is plausible that the physiological properties of these cells are also tuned to support the temporal precision demands of their local MSN circuit. Establishing correspondences and points of divergence between findings in mouse and primate will ultimately be essential for determining translational, clinical relevance of basal ganglia studies.

Multiple studies have shown phenotypic differences between D1 and D2 MSNs; specifically, D2 MSNs are more excitable than D1 MSNs and have a smaller dendritic area^57,58^. We found that while both D1 and D2 MSNs participated in the first, spatially-correlated axis of transcriptomic variation (E-LF-1 and M-LF-1), they were separated by the neuronal excitability-correlated E-LF-2, thus recapitulating these previously described differences; interestingly, though, we did not identify a morphology latent factor that was as aligned with MSN D1 / MSN D2 identity as E-LF-2. It is also possible that D1 and D2 MSNs in the same subregion of the striatum may be at somewhat different positions along the E-LF-1-aligned axis, leading to phenotypic differences deriving from that axis being attributed to D1 vs D2 identity. The simple D1 / D2 framework for MSNs has also been revised over the past decade by reports of a subpopulation of MSNs that co-expresses *Drd1* and *Drd2* (D1-Pcdh8^3^; eccentric SPNs^1^; D1H^4^). From a molecular standpoint, the D1 Sema5a Gaba subclass, which contains multiple T-types found in both CP and ACB, likely includes these hybrid MSNs^27^. We found that while these cells have a more similar molecular profile to D1 MSNs, with strong expression of *Drd1*, their highly excitable phenotype is more similar to (and in fact is on the extreme end of) D2 MSNs. Interestingly, *Tshz1*-expressing MSNs have been associated with populations exhibiting mixed D1/D2 molecular signatures, and D1 Sema5a cells express *Tshz1*, suggesting overlap between these populations^5,59^.

While continuous gradients structured much of the MSN diversity we observed, we also encountered subpopulations with more distinctive properties. Gene expression pattern and spatial location together suggested that the 0986 T-type may correspond to the patch-like *Htr7+* cells recently described in^4^. Similar to D1 Sema5a, these cells exhibited particularly high input resistance and low rheobase, even compared to other D2 MSNs, suggesting a highly excitable phenotype tuned to respond to weak inputs. These cells were also at the far D2 end of the gradient described by E-LF-2, suggesting they may in some sense represent an extreme version of the D2 phenotype. 0986 are members of a supertype with 2 other dorsal striatum T-types that also exhibit similar (albeit less extreme) physiological properties. Future work will be needed to understand how these unusual cells contribute to the striatal circuit.

Finally, we did not identify individual E-LFs or M-LFs that directly aligned with patch/matrix compartmentalization, another key axis of variation in MSNs. It is possible that cells in striosome or matrix compartments systematically differ from each other along dimensions that can be partially explained by multiple LFs together. Alternatively, while patch / matrix compartmentalization may have a strong gene expression difference, that variation may not be strongly coupled to the measured phenotypic features, though patch D1 MSNs have been observed to be more excitable than matrix D1 MSNs^33,34^.

### Striatal interneuron T-types

Recent work has proposed that striatal fast-spiking interneurons exist on a spatial continuum aligned with phenotype and *Pvalb* expression^7,60,61^. AP kinetics of these cells in our dataset varied continuously along a spatial gradient, with T-types 0833, 0834 and 0836 partitioning this space into discrete categories; these observations are in agreement with fast-spiking and fast-spiking-like cells described in the prior work, with 0836 corresponding to fast-spiking cells and 0833 corresponding to fast-spiking-like cells. Our results do not fully align with^61^ in the morphological space, as we only found one M-LF that appeared to capture morphological variation that was neither spatially linked nor distinguishing of the 3 T-types. However, morphological reconstructions from these cells were substantially more limited compared to our electrophysiology data, and our dataset spans a broader extent of the striatum due to our inclusion of ACB, which could possibly contribute to this discrepancy.

The striatal cholinergic interneuron population is not homogeneous; for example, differences in morphological properties have been observed across different striatal compartments^62^. Interestingly, we found that most cholinergic interneurons in our Patch-seq dataset mapped to the same T-type (0933); however, within this T-type we observed continuous variation in electrophysiological and morphological properties which also corresponded with location in the striatum. We note that the gene *Drd2* is one of the strong contributors to the gene expression gradient, as dopamine has been linked to changes in pause/rebound dynamics in cholinergic interneurons^63^.

### Cell types in other BG structures

We were able to use our multimodal dataset to map previously described cell types in the GPe, GPi, STN, SNr, SNc, and VTA to specific types within a whole-brain transcriptomic taxonomy. In GPe, we characterized the types corresponding to prototypic, arkypallidal, and cholinergic neurons through a combination of marker gene expression and physiological properties^27^. Similarly, in GPi we determined the specific transcriptomic identities in the taxonomy of the neuronal populations typically described by *Pvalb* and *Sst* expression and location within GPi (the core and the shell, respectively); the latter population are neurons that produce both GABA and glutamate^8^. Knowing the specific transcriptomic identifies of neurons in these structures can enable the development of specific genetic tools to access and perturb those sets of cells in targeted experiments^64^.

We also investigated the projection patterns of MSNs to these downstream BG cell types by integrating our transcriptomic-associated gradient to a whole-neuron morphology dataset through shared morphological features. Interestingly, it appears that this gradient is preserved to different extents in different structures (more so in GPi and GPe, less so in SNr), and certain cell types may preferentially get input from MSNs at different ends of the gradient. Future work using specific genetic tools may be able to probe these suggested connectivity patterns more directly.

### Limitations

There are many T-types across the BG, and our dataset does not represent all these types to the same extent, which limits the conclusions we can draw for undersampled T-types (e.g., cholinergic cells in the GPe; MSN-adjacent cell types in the OT D3 Folh and Chst9 subclasses). Our dataset was collected in a highly systematic way, which enables comparisons of multimodal properties that span all the cells; however, we may not be characterizing all the properties that differ across T-types with our standardized protocol. Our analysis of MSN projection patterns relies on comparisons of axon and dendrite locations in a common registered space, which cannot demonstrate actual connectivity, which may depend on differences at fine spatial scales or on cell-type specific rules. Finally, our findings here are observational and correlational; future studies that selectively perturb specific cell types are necessary to establish causal links between their multimodal features and circuit function.

## Conclusion

Patch-seq datasets such as the one presented here can bridge between large-scale, comprehensive transcriptomic taxonomies and atlases of cell types and other experimental modalities that provide insights into different aspects of brain circuit structure and function. For example, the procedure we used to link transcriptomic gradients to a whole-neuron morphology dataset via morphology could be applied to additional datasets in the future (such as large electron microscopy volumes). Integration of modalities ranging from gene expression to physiology to connectivity will be essential for building a comprehensive understanding of the cell type components of neural circuits.

## Methods

### Animal care and use

Mice were housed at ≤ 5 mice per cage and maintained on a twelve-hour light/dark cycle in a humidity- and temperature-controlled room. Water and food were available ad libitum. All experimental procedures that involved the use of mice were conducted with approved protocols in accordance with NIH guidelines and approved by the Allen Institute for Brain Science Institutional Animal Care and Use Committee (IACUC).

### Transgenic mice and labeling

Transgenic driver and reporter mice were used in some Patch-seq experiments to label populations of neurons for targeted recording. Characterizations of transgenic mouse line expression patterns are available in the Allen Institute for Brain Science Transgenic Characterization database (http://connectivity.brain-map.org/transgenic/search/basic)^65^. In other experiments, viral genetic tools were used to label subpopulations of cells^64,66–69^. Information about transgenic lines and viral tools can be found in Supplementary Data.

### Tissue processing and slicing procedure

To prepare acute brain slices for Patch-seq recording, adult female and male mice (95% aged P45–P70) were first fully anesthetized with 5% isoflurane inhalation. Intracardiac perfusion was performed with 25–50 mL of cold cutting artifical cerebrospinal fluid (ACSF; 0.5 mM calcium chloride (dihydrate), 25 mM D-glucose, 20 mM HEPES, 10 mM magnesium sulfate heptahydrate, 1.25 mM sodium phosphate monobasic monohydrate, 3 mM myo-inositol, 12 mM N-acetyl-L-cysteine, 96 mM N-methyl-D-glucamine chloride (NMDG-Cl), 2.5 mM potassium chloride, 25 mM sodium bicarbonate, 5 mM sodium L-ascorbate, 3 mM sodium pyruvate, 0.01 mM taurine, and 2 mM thiourea (pH 7.3)) which was continuously bubbling with a mixture of 95%*O*_2_/5%*CO*_2_. Sections were sliced on a vibrating microtome (Compresstome VF-300 vibrating microtome, Precisionary Instruments or VT1200S Vibratome, Leica Biosystems) at 350 µm thickness. To assist with registration to the Allen Mouse Common Coordinate Framework version 3 (CCFv3), a block-face image was collected before each section was cut (Mako G125B PoE camera with custom integrated software). Brain slices were placed in warm (34 °C) oxygenated cutting ACSF immediately after slicing for 10 minutes, then allowed to recover further in holding ACSF (2 mM calcium chloride (dihydrate), 25 mM D-glucose, 20 mM HEPES buffer, 2 mM magnesium sulfate heptahydrate, 1.25 mM sodium phosphate monobasic monohydrate, 3 mM myo-inositol, 12.3 mM N-acetyl-L-cysteine, 84 mM sodium chloride, 2.5 mM potassium chloride, 25 mM sodium bicarbonate, 5 mM sodium L-ascorbate, 3 mM sodium pyruvate, 0.01 mM taurine, and 1 mM thiourea (pH 7.3)), bubbling with a mixture of 95%*O*_2_/5%*CO*_2_ at room temperature until transferred to the microscope for recording.

### Patch-clamp recording

Patch-seq recordings were performed as previously described^11,28,29^. Slices were bathed in warmed (34 °C) recording ACSF (2 mM calcium chloride (dehydrate), 12.5 mM D-glucose, 1 mM magnesium sulfate heptahydrate, 1.25 mM sodium phosphate monobasic monohydrate, 2.5 mM potassium chloride, 26 mM sodium bicarbonate, and 126 mM sodium chloride (pH 7.3)) and continuously bubbled with 95% O_2_/5% CO_2_. The bath solution contained blockers of fast glutamatergic (1 mM kynurenic acid) and GABAergic synaptic transmission (0.1 mM picrotoxin). Thick-walled borosilicate glass (Warner Instruments, G150F-3) electrodes were pulled (Narishige PC-10) with a resistance of 4–5 MΩ.The electrodes were filled with ∼1.0 to 1.5 µL of internal solution with biocytin (110 mM potassium gluconate, 10.0 mM HEPES, 0.2 mM ethylene glycol-bis (2-aminoethylether)-N,N,N^′^,N^′^-tetraacetic acid, 4 mM potassium chloride, 0.3 mM guanosine 5^′^-triphosphate sodium salt hydrate, 10 mM phosphocreatine disodium salt hydrate, 1 mM adenosine 5^′^-triphosphate magnesium salt, 20 µg/mL glycogen, 0.5U/µL RNAse inhibitor (Takara, 2313A) and 0.5% biocytin (Sigma B4261), pH 7.3). The pipette was mounted on a Multiclamp 700B amplifier headstage (Molecular Devices) attached to a micromanipulator (PatchStar, Scientifica).

Electrophysiology signals were recorded with an ITC-18 Data Acquisition Interface (HEKA). The software package MIES (https://github.com/AllenInstitute/ MIES/), written in Igor Pro (Wavemetrics), was used to generate command currents/potentials, process recorded signals, and collect amplifier metadata.

Data were filtered (Bessel) at 10 kHz and digitized at 50 kHz. Current-clamp data are reported uncorrected for the measured liquid junction potential^70^ of −14 mV between the electrode and bath solutions. Before data collection, all surfaces, equipment, and materials were cleaned thoroughly in the following manner: wiped down with DNA away (Thermo Scientific), RNAse Zap (Sigma-Aldrich), and finally with nuclease-free water.

After formation of a stable seal and whole-cell break-in, the resting membrane potential of the neuron was recorded (typically within the first minute). A bias current was injected, either manually or automatically using algorithms within the MIES data acquisition package, for the remainder of the experiment to maintain that initial resting membrane potential. Bias currents remained stable for a minimum of 1 s before each stimulus current injection. If cells fired action potentials spontaneously without current injection, additional hyperpolarizing current was injected to suppress the spontaneous activity. In a subset of experiments, spontaneous activity was monitored for 60 seconds in tight-seal cell-attached configuration in voltage clamp mode, prior to breaking into the cell.

To be included in electrophysiological analyses, neurons needed to fulfill several criteria; a *>*1 GΩ seal recorded before break-in, an initial access resistance *<*20 MΩ and *<*20% of the *R*_input_. For an individual sweep to be included, the following criteria were applied: (1) the bridge balance was *<*20 MΩ and *<*20% of *R*_input_; (2) bias (leak) current within ±100 pA; and (3) root mean square noise measurements in a short window (1.5 ms, to gauge high frequency noise) and longer window (500 ms, to measure patch instability) were *<*0.07 mV and *<*0.5 mV, respectively.

After electrophysiological recording was completed, the pipette was centered on the soma or placed near the nucleus, if visible. A small amount of negative pressure (∼ −30 mbar) was applied to begin cytosol extraction and to attract the nucleus to the tip of pipette. After approximately one minute, the soma visibly shrank and/or the nucleus was observed near the tip of the pipette. With maintained negative pressure, the pipette was retracted; slow, continuous movement was maintained while monitoring the pipette seal. Once the pipette seal reached *>*1 GΩ and the nucleus was visible at the tip of the pipette, the speed was increased to remove the pipette from the slice. The pipette containing internal solution, cytosol, and the nucleus was removed from the pipette holder, and its contents were expelled into a PCR tube containing the lysis buffer (Takara, 634894).

Metadata for all Patch-seq neurons including in this study are located in Supplementary Data.

### Macaque Patch-seq data

The macaque Patch-seq cells used here were first collected by Liu *et al.* [37]. Detailed experimental procedures are described in that study. In most cases, those Patch-seq recordings were performed in the same manner as the mouse recordings described in this study. Data from 437 macaque MSNs were analyzed using the same methods and software (described below). Non-human primate brain specimens were obtained from animals designated for the Washington National Biomedical Research Center’s (WaNBRC) Tissue Distribution Program, and all procedures involving non-human primates were approved by the University of Washington’s Institutional Care and Use Committee (IACUC) and conformed to the NIH’s Guide for the Care and Use of Laboratory Animals.

### Electrophysiology feature analysis

Electrophysiological features were measured from responses elicited by short (3 ms) current pulses and long (1 s) current steps as previously described^28,29,71^. Action potentials (APs) were detected, and the threshold, peak, fast trough, and width (at half-height) were calculated for each AP, along with the ratio of the peak upstroke dV/dt to the peak downstroke dV/dt (upstroke/downstroke ratio). Several voltage trajectories (the initial AP elicited by the lowest-amplitude current pulses and steps, the derivatives of those APs, and the interspike interval) were also extracted for analysis. AP features across responses to long current steps were averaged in time bins and concatenated across step amplitudes; bins without APs contained interpolated values from their neighbors. This was done for steps starting at a given cell’s rheobase and increasing at 10 pA intervals. Sweeps from intervals without data were interpolated from sweeps at neighboring intervals. Subthreshold responses to hyperpolarizing current steps were analyzed as before by downsampling to 10 ms bins and concatenating responses from different stimulus amplitudes (ranging from -90 pA to -10 pA). Sparse principal component analysis was performed separately on data from each of these categories (e.g., AP waveform, AP features across current steps), and sparse principal components (sPCs) that exceeded 1% adjusted explained variance were kept. This yielded 51 sPCs in total from twelve data categories. The components were z-scored and combined to form a reduced dimension electrophysiology feature matrix. Additional features such as the membrane time constant, input resistance, and hyperpolarization-induced sag were calculated from responses to long hyperpolarizing current steps as previously described^71^.

### cDNA amplification and library construction

Using cellular material collected from Patch-seq experiments, nuclear and cytosolic mRNA was reverse transcribed and the resulting cDNA was sequenced using the SMART-Seq v4 method described in Tasic *et al.* [72]. The SMART-Seq v4 Ultra Low Input RNA Kit for Sequencing (Takara, 634894) was used to reverse transcribe poly(A) RNA and amplify full-length cDNA according to the manufacturer’s instructions. Reverse transcription and cDNA amplification was performed for 20 PCR cycles in 0.65 mL tubes, in sets of 88 tubes at a time. At least 1 control 8-strip was used per amplification set, which contained 4 wells without cells and 4 wells with 10 pg control RNA. Control RNA was either Mouse Whole Brain Total RNA (Zyagen, MR-201) or control RNA provided in the SMART- Seq v4 kit. All samples proceeded through Nextera XT DNA Library Preparation (Illumina FC-131-1096) using either Nextera XT Index Kit V2 Set A-D (FC-131-2001,2002,2003,2004) or custom dual-indexes provided by IDT (Integrated DNA Technologies). Nextera XT DNA Library prep was performed according to manufacturer’s instructions except that the volumes of all reagents including cDNA input were decreased either to 0.4x or to 0.2× by volume. Each sample was sequenced to approximately 500,000 - 1 million reads.

### Sequencing data processing

Fifty-base-pair paired-end reads were aligned to mm10 GENCODE vM23/Ensembl 98 reference genome, downloaded from 10X cell ranger (refdata-cellranger-arc-mm10-2020-A-2.0.0). Sequence alignment was performed using STAR aligner (v2.7.1a) with default settings. PCR duplicates were masked and removed using STAR option “bamRemoveDuplicates.” Only uniquely aligned reads were used for gene quantification. Gene counts were computed using the R Genomic

Alignments package^73^ summarizeOverlaps function using “IntersectionNotEmpty” mode for exonic and intronic regions separately. Exonic and intronic reads were added to calculate total gene counts. Data were analyzed as counts per million reads (CPM).

### Transcriptomic mapping and analysis

The reference dissociated cell single-cell transcriptomics data and MERFISH data used in this study were from the whole mouse brain cell atlas^2^. Patch-seq cells were mapped to the whole mouse brain taxonomy^2^ using the Hierarchical Approximate Nearest Neighbor (HANN) method implemented in the scrattch-mapping package (https://github.com/alleninstitute/scrattch-mapping). This method involved traversing the taxonomy hierarchy, selecting offspring node-differentiating marker genes at each node, and finding the approximate nearest neighbor T-type using marker gene correlation as the distance metric.

As in previous work^28,29^, we evaluated the T-type mappings by considering the confidence with which a Patch-seq transcriptome mapped to one or more reference T-types, and the expected level of ambiguity between reference T-types. We classified mapping quality measures (based on differentially expressed gene correlation and the Kullback-Leibler (KL) divergence between the mapping probability distributions of Patch-seq cells and the reference mapping probability distribution) into ”highly consistent”, ”moderately consistent” and ”inconsistent” categories. Cells with ”inconsistent” mapping were not included in further analyses. We imposed an additional requirement for minimum number of genes detected (2000).

### Morphological reconstruction

Neuronal morphologies were imaged and reconstructed as previously described^28,29,71^. Neurons were filled with biocytin via the patch pipette. To visualize the label, a horseradish peroxidase (HRP) enzyme reaction using diaminobenzidine (DAB) as the chromogen was used afterwards. 4,6-diamidino-2-phenylindole (DAPI) stain was also used to help visualize cytoarchitectural features and landmarks.

Slices from Patch-seq experiments were mounted on slides and imaged as described previously^71^. Images were collected with an upright AxioImager Z2 microscope (Zeiss, Germany) equipped with an Axiocam 506 monochrome camera and 0.63x optivar lens. Two-dimensional tiled overview images were also captured (Zeiss Plan-NEOFLUAR 20X/0.5) in brightfield transmission and fluorescence channels. Higher resolution image stacks of individual cells were acquired in the transmission channel only for the purpose of morphological reconstruction. Light was transmitted using an oil-immersion condenser (1.4 NA). High-resolution, multi-tile image stacks were captured (Zeiss Plan-Apochromat 63x/1.4 Oil or Zeiss LD LCI Plan-Apochromat 63x/1.2 Imm Corr) at an interval of 0.28 µm (1.4 NA objective) or 0.44 µm (1.2 NA objective) along the Z axis.

Image tiles were stitched in ZEN software and exported as single-plane TIFF files. Anatomical locations of recorded neurons were determined based on DAPI stained overview images described above. The soma position of reconstructed neurons were recorded for subsequent analysis. Individual cells were manually aligned to the CCFv3 by matching the overview image of the slice with a “virtual” slice at an appropriate location and orientation within the CCFv3.

Dendritic reconstructions were completed for a subset of neurons with high quality transcriptomics, electrophysiology, and labeling. Reconstructions were based on the 63X image stacks described above. Stacks were run through a Vaa3D-based image processing and reconstruction pipeline^74^. An automated reconstruction of the neuron was produced using TReMAP^75^. Alternatively, initial reconstructions were created manually using the reconstruction software PyKNOSSOS (Ariadne-service) or through the citizen neuroscience game Mozak (Mozak.science)^76^. Automated or manually initiated reconstructions were then extensively manually corrected and extended using a range of tools (e.g., virtual finger, polyline) in the Mozak extension (Zoran Popovic, Center for Game Science, University of Washington) of Terafly tools^77,78^ in Vaa3D. Where possible, the local axon was also reconstructed. Additionally, all neurons eligible for reconstruction were automatically segmented and post-processed to produce a quantifiable neuron reconstruction using the approach described in Gliko *et al.* [79].

After 3D reconstruction, morphological features were calculated using similar methods as previously described^28,29,71^. Because the basal ganglia neurons are not in a laminar structure (unlike the previous studies in visual cortex), neuron morphologies were aligned to the CCFv3 and measurements such as the extent and bias of dendritic arbors were calculated with respect to global anatomical directions (i.e., anterior/posterior, medial/lateral, dorsal/ventral). Other features, such as total dendritic length, number of branches, etc., did not depend on neuron location or orientation. In addition, Sholl analysis was performed to quantify dendritic arbor intersections in a series of concentric shells for each neuron, followed by principal components analysis on the set of Sholl profiles to calculate additional morphological features (Sholl (PC1), etc.).

### MET-type definition

MET-types were defined for basal ganglia neurons that had data for all three data modalities (morphological, electrophysiological, and transcriptomic) using methods previously used for cortical neurons^28,29^. Electrophysiological and morphological features were used first to define ME-clusters by several clustering methods; consensus clusters were identified based on the combined results^28,29,71^. Next, a graph was constructed from nodes that represented either cells or ME-type/T-type combinations. Edges connected cell nodes to ME-/T-type nodes with edge weights equal to the cells’ T-type mapping probabilities (from the original T-type mapping procedure) and ME-cluster mapping probability (from subsampled random forest classification). Cells were also connected to each other by the average of their pairwise correlation across all three modalities; only the top 1%, 1.5%, or 2% of correlation edges were used.

The Leiden community detection algorithm^80^ was used to group strongly-connected nodes into MET-types (Extended Data Fig. 1). The procedure was repeated with twenty different random seeds for each of the three edge weight cut-offs. Final MET-types were defined from consensus clustering (as with ME-types) from across the sixty total runs. Cells that did not reliable co-cluster with other cells in the consensus cluster (*<* 50% average co-clustering rate) were not given a final MET-type assignment (*n* = 29). This procedure resulted in 163 cells being assigned to 23 MET-types.

### Sparse reduced-rank regression

To identify the strongest correspondences between transcriptomic expression patterns and electrophysiological and morphological characteristics of Patch-seq neurons, we performed sparse reduced-rank regression^30,81^. In this method, a multivariate regression is performed to predict either electrophysiological or morphological features from gene expression data using a small number of latent factors as an intermediate layer (reduced-rank regression), combined with an elastic net regularization to select a sparse set of contributing genes and to constrain the model weights^30^.

For this study, we used our previously described re-implementation^29^ of the original sparse reduced-rank regression (sparse RRR) Python code (https://github.com/berenslab/patch-seq-rrr in the R language using the glmnet package^82^ for improved performance when selecting hyperparameters and integration with other aspects of our code base. We performed sparse RRR separately for striatal medium spiny neurons, striatal cholinergic interneurons, and striatal fast-spiking interneurons. For the fits, highly-variable genes were first selected for each group using Brennecke’s method (https://github.com/ AllenInstitute/scrattch.hicat/). Features to be fit in the multivariate sparse RRR were selected by first performing elastic net fits based on gene expression on individual features; features that could be fit with a cross-validated *R*^2^ *>* 0.1 were kept for sparse RRR. Next, cross-validation was used to select the optimal rank (i.e., number of latent factors), alpha (parameter controlling the trade-off between ridge and lasso penalties), and lambda (overall penalty strength) hyperparameters for each group and modality (see Extended Data Figure 5). For each subclass and modality, we also held out 15% of the cells as a test dataset that was not a part of hyperparameter selection or final regression fitting.

After performing sparse RRR with the selected hyperparameters, we visualized the results using side-by-side bi-plots^30,81^ with a few selected highly correlated genes and features (see Figures 2, 5 and Extended Data Figure 6). The electrophysiological sparse RRR was performed on the electrophysiology sPCs (see above), but we also calculated correlations between latent factors and traditionally defined electrophysiology features (e.g., mean AP width) for ease of interpretation. To estimate latent factors for WNM neurons (see below), we used the sparse RRR weights on the matched morphological feature set (as when calculating the morphology side of the paired bi-plots using Patch-seq data).

### Patch/matrix score estimation

To assess the feature relationships with patch/matrix membership, an elastic net regression model was trained to predict a previously described patch/matrix score, defined as log(*Kremen1*) + log(*Sema5b*) - log(*Id4*)^4^ using reference dissociated scRNA-seq MSNs and a broader set of highly variable genes (*R*^2^ = 0.64). This enabled robust estimation of a patch/matrix score across different datasets and reduced the sensitivity to noise in individual gene measurements.

### Whole neuron morphology reconstructions

The whole neuron morphologies of MSNs were part of the previously published study by Peng *et al.* [54]. Neurons were reconstructed from GFP-labeled brains and imaged using the fMOST technique; the details of the experimental procedures are described in the referenced study. The morphological features of the dendritic arbors of those cells were calculated in the same way as for the Patch-seq neuron reconstructions. However, there are systematic differences in the morphological features due to technical differences in the techniques (e.g., Patch-seq neurons are contained within 350 µm thick slices. To align morphological feature distributions between datasets, we employed a procedure analogous to that described in Sorensen *et al.* [29], with one modification necessitated by the absence of laminar depth information in basal ganglia tissue. In the original approach, point clouds were constructed using soma depth as a fixed reference axis, against which individual morphological features were aligned via a linear transformation minimizing Chamfer distance. Because basal ganglia nuclei lack the laminar organization that makes cortical soma depth a meaningful ordering variable, we instead used morphological principal component 1 (PC-1), computed from the full dendritic feature set prior to alignment, as a proxy for the static axis. As in the original method, only the feature-value component of each point was subject to transformation; the reference axis was held fixed. This alignment was performed independently for each morphological feature. The use of PC-1 as a proxy preserves the intent of the original method — anchoring feature alignment to a cell-intrinsic ordering variable that captures systematic variation across the population — while remaining applicable to non-laminated structures.

We assigned CP subdivisions to fMOST cells based on soma location and CCFv3 masks^22^. We quantified the convergence of axonal inputs from fMOST cells arising from CP subdivisions by constructing voxelized representations of axon terminal and branch node density within each target structure, and accumulating voxel-wise counts of axons. To account for differences in sampling across the 4 CP subdivisions, we normalized voxel counts by the number of cells from each subdivision that contributed axons to the target structure. The resulting maps were smoothed with a Gaussian kernel (sigma = 75 µm). We defined binary ‘presence’ of a CP subdivision at each voxel if their smoothed density exceeded a given threshold (20% of the maximum value for that subdivision in the structur). This allowed us to quantify the number of CP subdivisions contributing to each voxel. To quantify variation in predicted M-LF-1 values in target structures, we aggregated voxels into 30 µm × 30 µm × 30 µm bins, and averaged M-LF-1 values of contributing cells to each bin (each cell counting at most once). For visualization, density and subdivision convergence maps were projected into coronal space by taking the maximum value, and restricted to the boundaries of the structure. Mean M-LF-1 values were projected in coronal space by taking the mean value. To quantify M-LF-1 distribution in fMOST axons in the local milieu of SNr, GPi and GPe T-types, we first fit a radius-based nearest neighbor model on all axon terminals and branches of fMOST reconstructions in a given structure, thus enabling identification of axons within 50 µm of dendrites from Patch-seq reconstructions. For each Patch-seq cell, we aggregated unique axon neighbors from the fMOST dataset and summarized the distribution of their M-LF-1 values, averaging across cells within each T-type.

### Statistics and research design

No statistical methods were used to predetermine sample sizes, but sample sizes here are similar to those reported in previous publications. No randomization was used during data collection as there was a single experimental condition for all acquired data. The different stimulus protocols were not presented in a randomized order. Data collection and analyses were not performed blind to the conditions of the experiments as there was a single experimental condition for all acquired data.

Correlations were measured by the nonparametric Spearman rank correlation coefficient unless otherwise noted. Differences across multiple groups were assessed using one-way analysis of variance (ANOVA), with post-hoc pairwise comparisons using Tukey’s test to correct for multiple comparisons. Comparisons between two groups were performed using Welch’s two-sample t-test; for multiple features, p-values were corrected for multiple comparisons using the Bonferroni method. Statistical significance was defined as p*<* 0.05.

Data are reported as mean ± standard deviation and averaged AP waveforms, AP phase plots and subthreshold voltage responses are visualized as mean ± standard error unless otherwise noted.

### Data and software availability

Transcriptomic data supporting the findings of this study will be made available at the NeMO archive. Similarly, electrophysiological data for this study will be available at the DANDI archive, and morphological reconstructions from this study will be available at the BIL archive.

The electrophysiology data acquisition software (MIES) used for this study is available at https://github.com/alleninstitute/mies. The morphological reconstruction software (Vaa3D-TeraFLY-Mozak is freely available at http://home.penglab.com/proj/vaa3d/home/index.html and the code is available at https://github.com/Vaa3D. The code for electrophysiological and morphological feature analysis and clustering is available as part of the open-source Allen SDK repository (https://github.com/AllenInstitute/AllenSDK), skeleton-keys repository (https://skeleton-keys.readthedocs.io/en/latest/), py-ropractor repository (https://github.com/AllenInstitute/pyropractor/), IPFX repository (https://github.com/alleninstitute/ipfx), and DRCME repository (https://github.com/alleninstitute/drcme). The code for processing and quantifying WNM reconstructions is part of the morph-utils repository (https://github.com/matthewMallory/morph_utils). The re-implementation of sparse reduced-rank regression is available in the repository of a previous study (https://github.com/AllenInstitute/exc_vis_manuscript).

## Supporting information

Patch-seq sample metadata

Transgenic and viral tools

## Acknowledgments

We are grateful to the Transgenic Colony Management, Neurosurgery & Behavior, Lab Animal Services, Molecular Biology, Histology, Viral Technology, and Imaging teams at the Allen Institute for technical support. We thank Ximena Opitz-Araya and Refugio Martinez for AAV vector molecular cloning support. The research was funded in part by National Institutes of Health (NIH) grants U01NS132267 (S.A.S. and T.J.), UM1MH130981 (E.S.L. and H.Z.), and UF1MH128339 (B.T. and J.T.T.). The WaNBRC is supported by the NIH Office of Research Infrastructure Programs (ORIP) under awards P51OD010425 and U420D011123. The content is solely the responsibility of the authors and does not necessarily represent the official views of NIH and its subsidiary institutes. This work was also supported by the Allen Institute for Brain Science. We thank Allan Jones and Rui Costa for leadership and guidance. We wish to thank Allen Institute founders, P. G. Allen and J. Allen, for their vision, encouragement and support.

## Author Contributions

**Conceptualization**: A.Bu., E.S.L., J.T.T., B.T., Z.Y., H.Z., B.K., T.J., S.A.S., B.R.L., N.W.G. **Methodology**: A.Bu., M.Ma., R.D., C.L., X.C., X.L., S.W., A.Bh., S.D., Z.C.J., M.Mc., L.P., S.T.R., J.Ar., K.A.S., Q.W., J.T.T., Z.Y., T.J., S.A.S., B.R.L., N.W.G. **Software**: A.Bu., M.Ma., C.L., X.C., S.W., N.J., S.D., Z.Y., T.J., N.W.G. **Validation**: A.Bu., B.R.L., N.W.G. **Investigation**: R.D., R.Ma., L.A., J.An., A.A., K.Ba., S.B., Z.B., D.B., K.Bl., P.B., K.Br., T.Car., T.Cas., N.I.D., T.E., R.E., A.G., J.G., K.H., Z.C.J., M.K., G.L., J.M., A.Ma., R.Mc., A.Mc., M.Mc., K.N., L.N., A.O., C.A.P., L.P., R.R., S.T.R., I.R., C.R., A.R., M.T., J.T., S.V., M.C.V., J.Ar., N.D., S.D.H. **Resources**: R.D., N.J., E.A., A.Bh., L.P., K.R., S.D.H., K.A.S., Q.W., J.W., B.T., S.A.S., B.R.L. **Data Curation**: A.Bu., M.Ma., R.D., X.C., X.L., S.W., J.G., M.T., K.A.S., B.R.L., N.W.G. **Writing - Original Draft**: A.Bu., N.W.G. **Writing - Review & Editing**: A.Bu., M.Ma., R.D., X.L., Q.W., J.T.T., H.Z., B.K., S.A.S., B.R.L., N.W.G. **Visualization**: A.Bu., N.W.G. **Supervision**: R.D., M.Mc., L.P., K.R., J.Ar., N.D., L.E., S.D.H., K.A.S., S.M.S., J.W., E.S.L., J.T.T., B.T., Z.Y., H.Z., B.K., T.J., S.A.S., B.R.L., N.W.G. **Project Administration**: R.D., L.E., K.G., L.K., S.M.S., B.R.L. **Funding Acquisition**: E.S.L., J.T.T., B.T., H.Z., T.J., S.A.S.

---

**Extended Data Figure 1:**
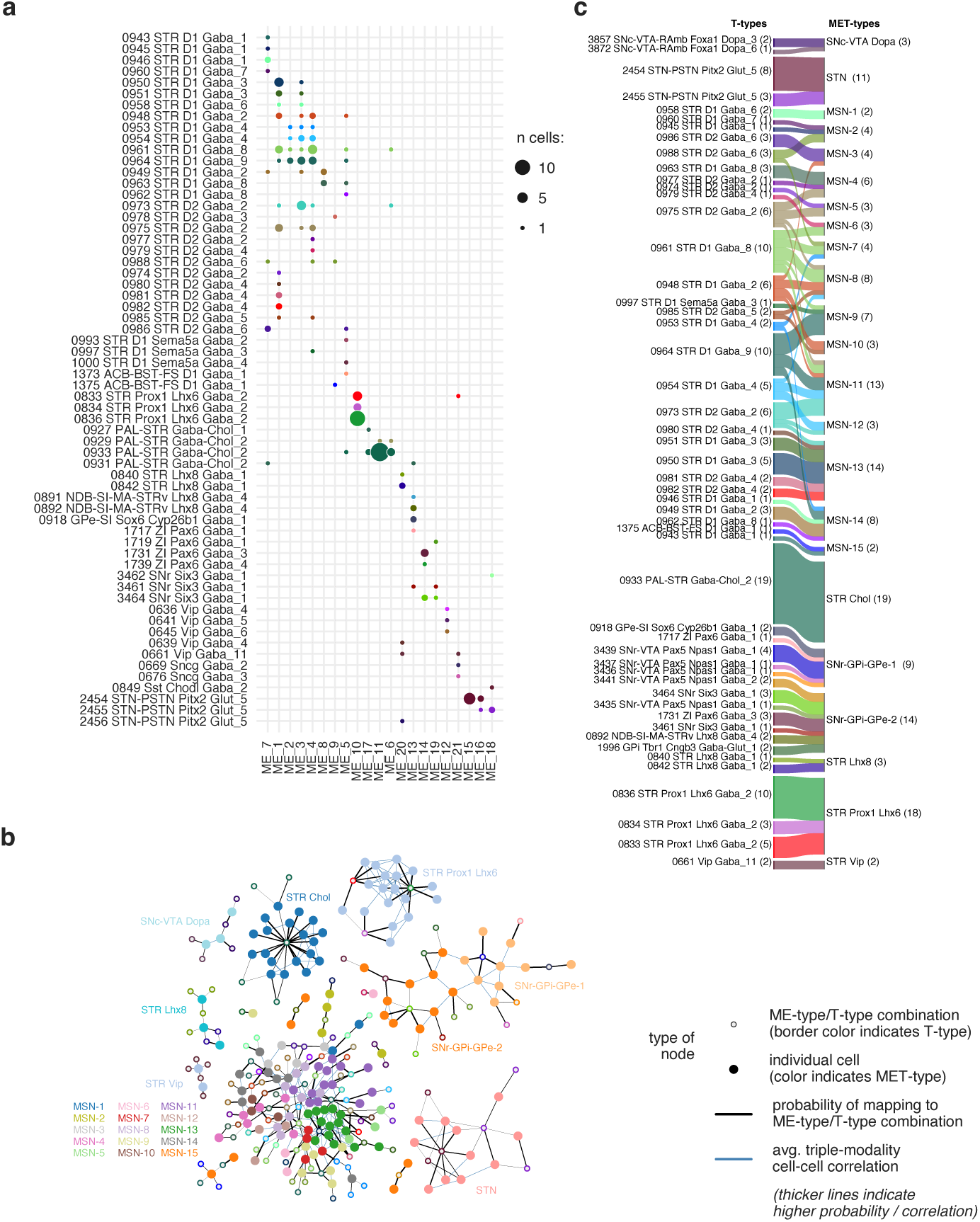
Definition of basal ganglia MET-types. **a**, Relationship between T-types and ME-types (*n* = 192 cells). Dot size indicates the number of cells with a given ME-type/T-type combination. **b**, Community-detection grouping of cells into MET-types (colors of solid nodes). Solid nodes are individual cells; hollow nodes are ME-/T-type combinations (as in (**a**)). Black edges represent mapping probabilities of cells to specific ME-/T-type combinations; blue edges represent average triple-modality cell-cell correlations. Thicker lines indicate higher probabilities and correlations. Only cell-cell correlations within the top 1% of strength values are shown. Node positions were determined by a spring model layout. **c**, Relationship between T-types (left) and MET-types (right).

**Extended Data Figure 2:**
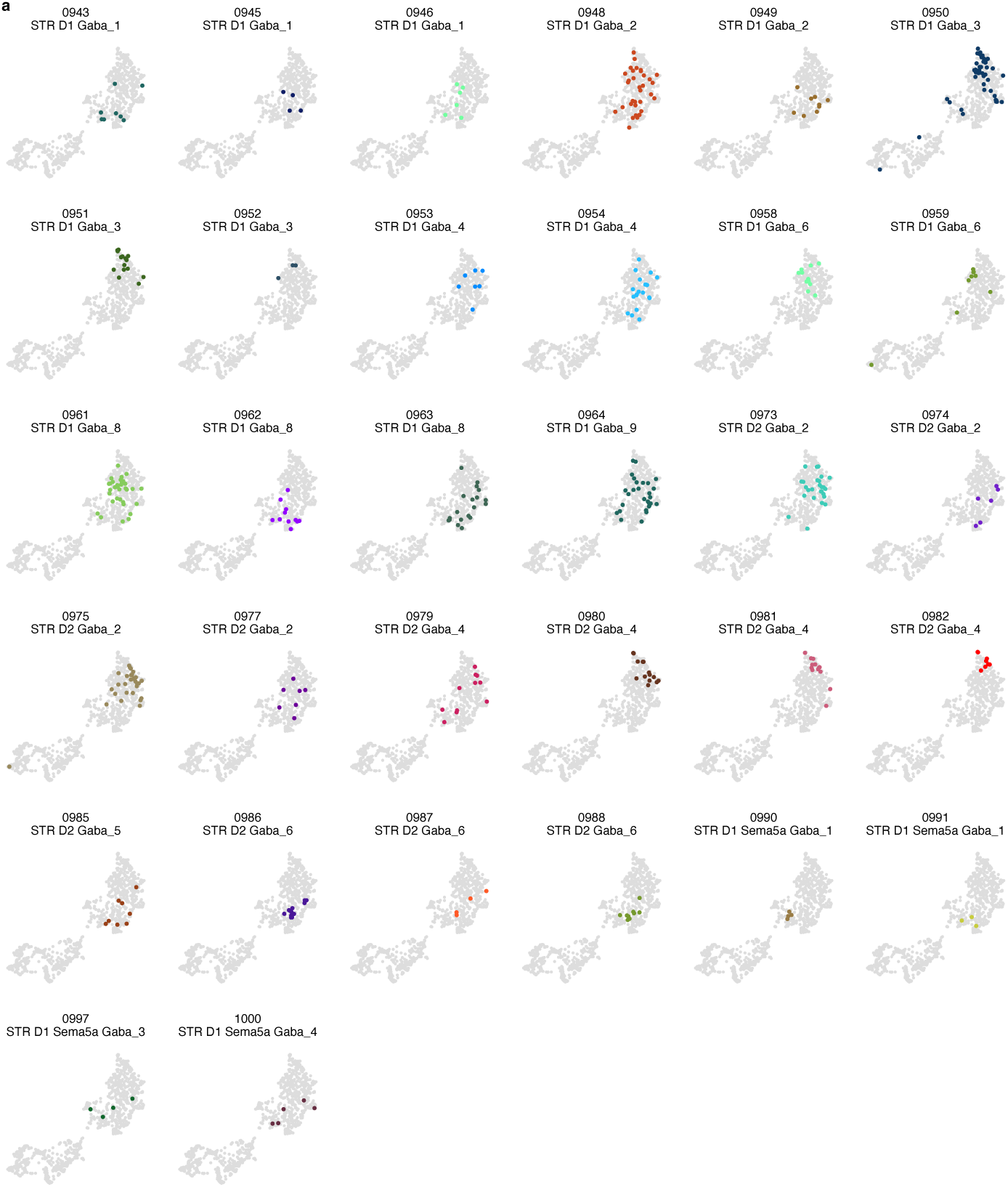
Electrophysiology UMAPs of MSN T-types. **a**, Electrophysiology UMAP highlighting MSN T-types with a minimum of 3 cells.

**Extended Data Figure 3:**
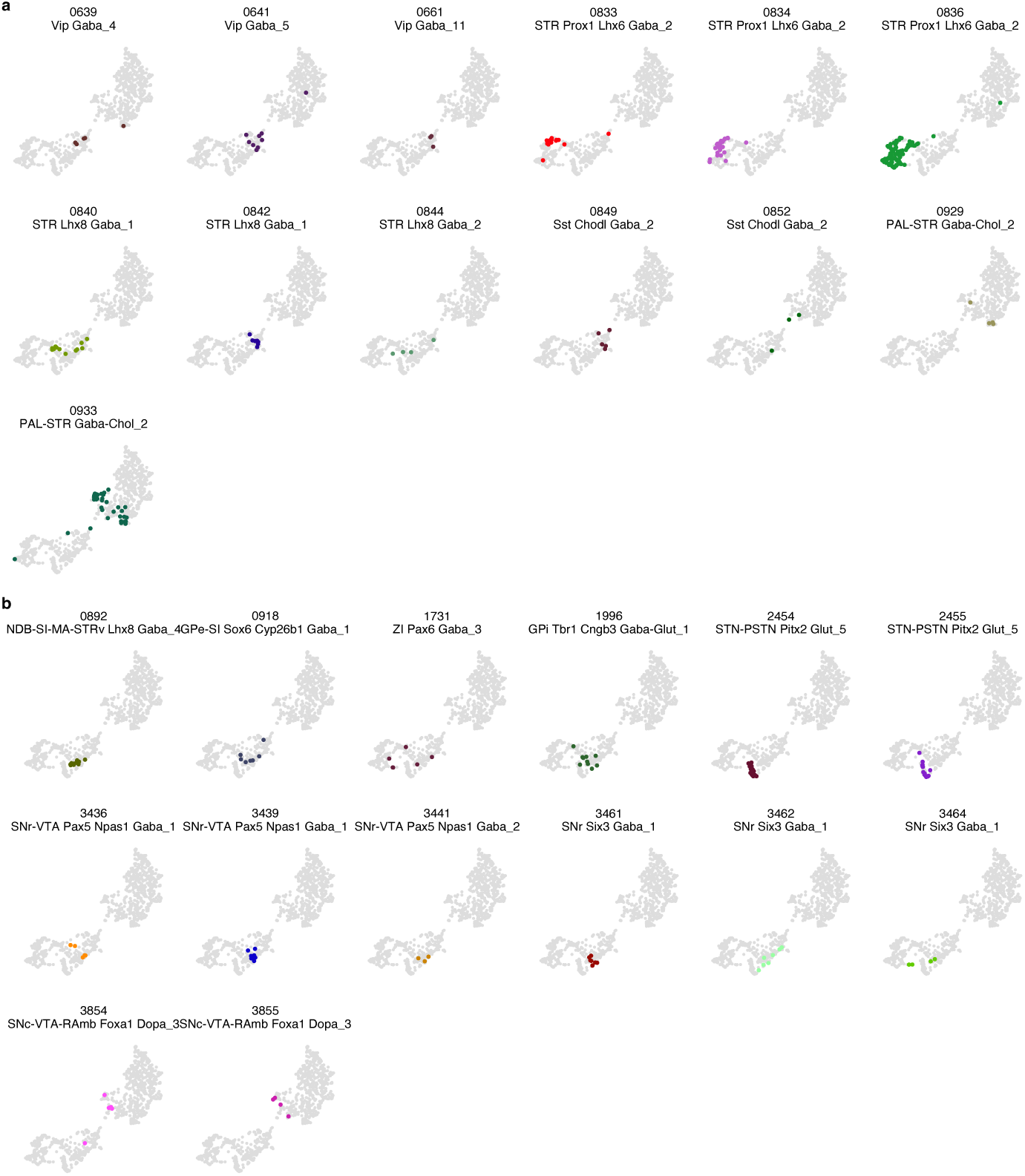
Electrophysiology UMAPs of other basal ganglia T-types. **a**, Electrophysiology UMAP highlighting striatal interneuron T-types with a minimum of 3 cells. **b**, Electrophysiology UMAP highlighting GPe, GPi, STN, SNr and SNc/VTA T-types with a minimum of 3 cells.

**Extended Data Figure 4:**
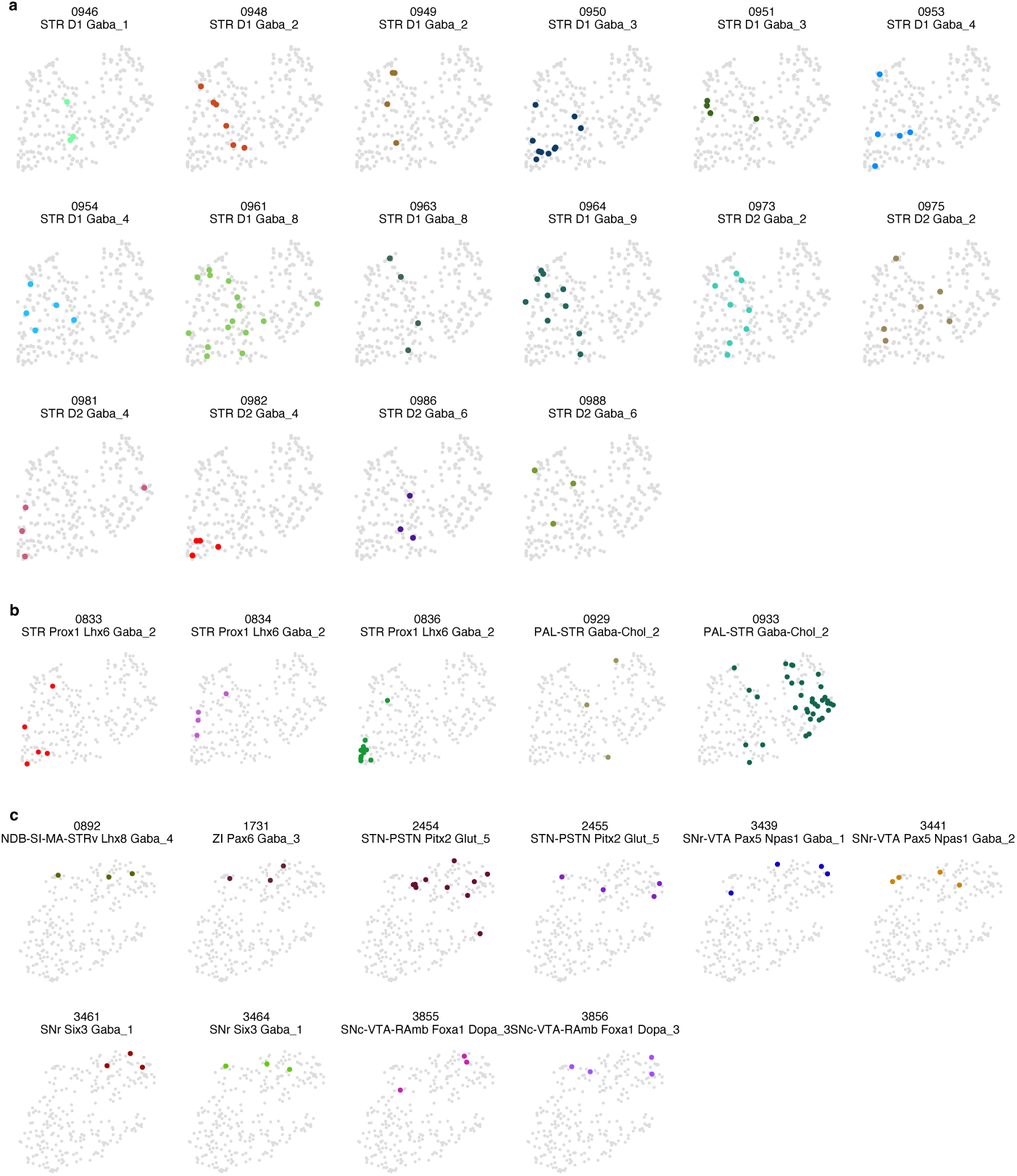
Morphology UMAPs of basal ganglia T-types. **b**, Morphology UMAP highlighting MSN T-types with a minimum of 3 cells. **b**, Morphology UMAP highlighting striatal interneuron T-types with a minimum of 3 cells. **c**, Morphology UMAP highlighting GPe, GPi, STN, SNr and SNc/VTA T-types with a minimum of 3 cells.

**Extended Data Figure 5:**
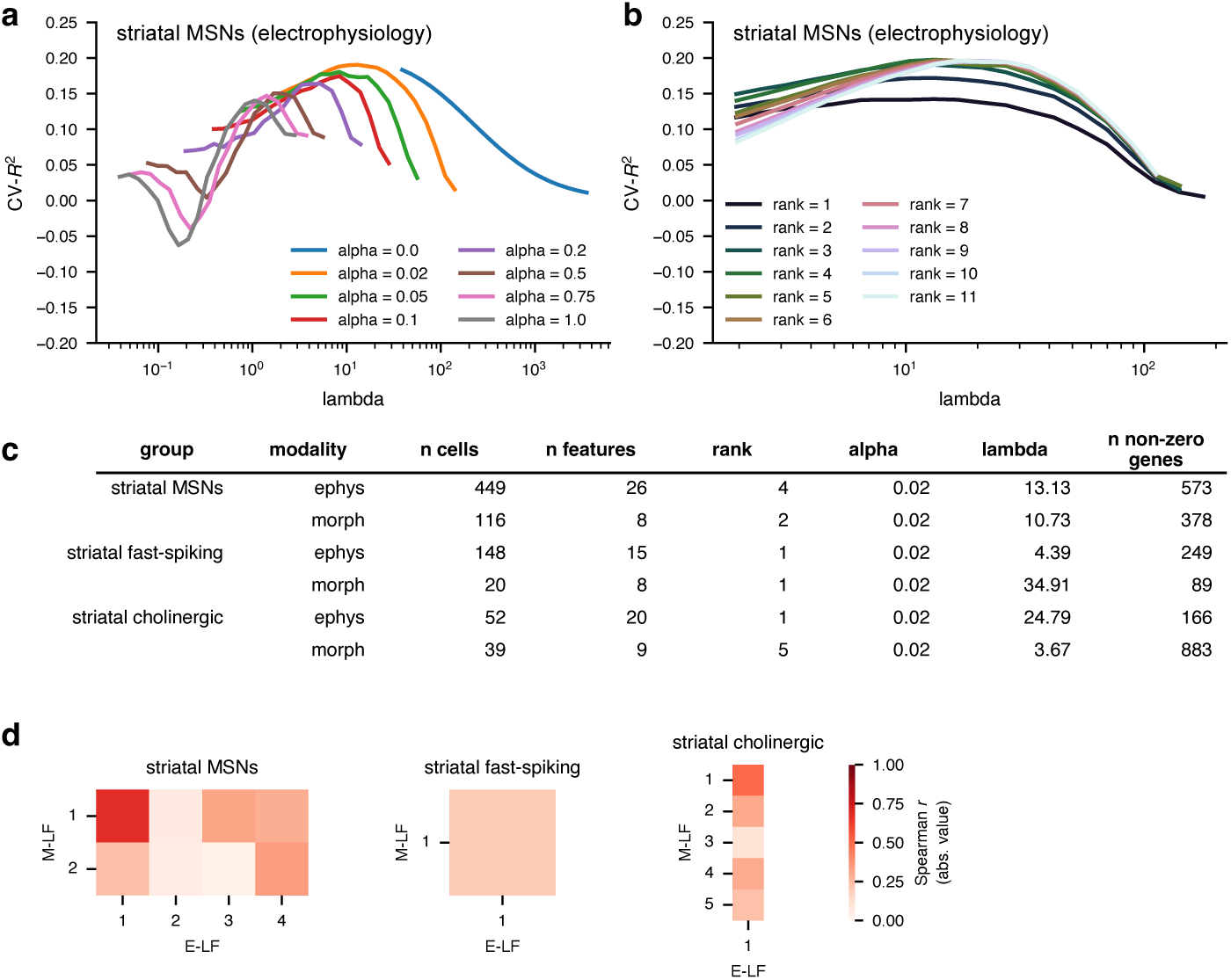
Sparse reduced-rank regression hyperparameter selection and cross-modality comparison. **a**, Selection of the alpha hyperparameter by cross-validation for striatal MSNs electrophysiology features. **b**, Selection of the rank hyperparameter by cross-validation for striatal MSNs electrophysiology features. **c**, Hyperparameters and other sRRR characteristics for each cell type group and modality. **d**, Spearman correlations between electrophysiology and morphology latent factors (calculated from gene expression) by cell type group (absolute value of the correlation is shown since the relative signs of the latent factors are arbitrary).

**Extended Data Figure 6:**
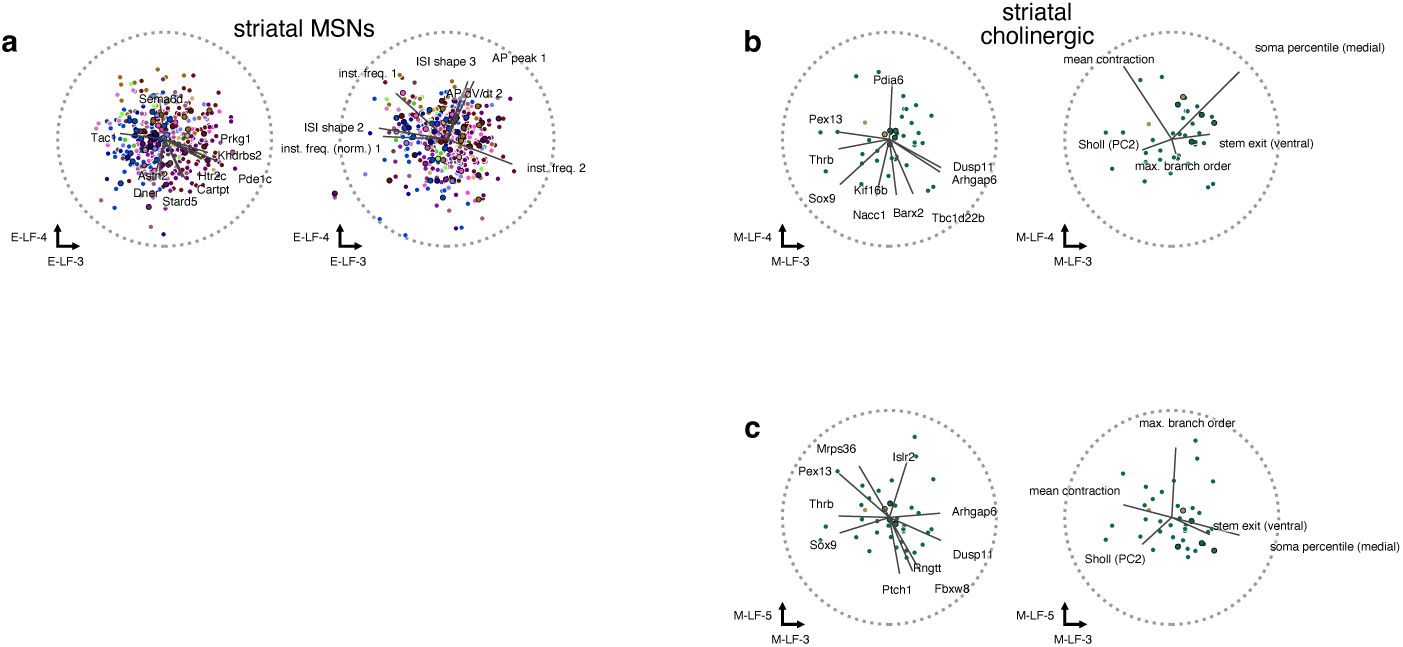
**Other latent factors from sparse reduced-rank regression. a-c**, Bi-plots of other latent factors from striatal MSNs (**a**) and striatal cholinergic interneurons (**b-c**). Latent factors are calculated from genes (left) and features (right) for electrophysiology (E-LFs) and morphology (M-LFs). Genes and features with the highest correlations with latent factors are shown (lines); dotted circle indicates correlation equal to 1. Cells that were part of a held-out test dataset are marked with black borders.

**Extended Data Figure 7:**
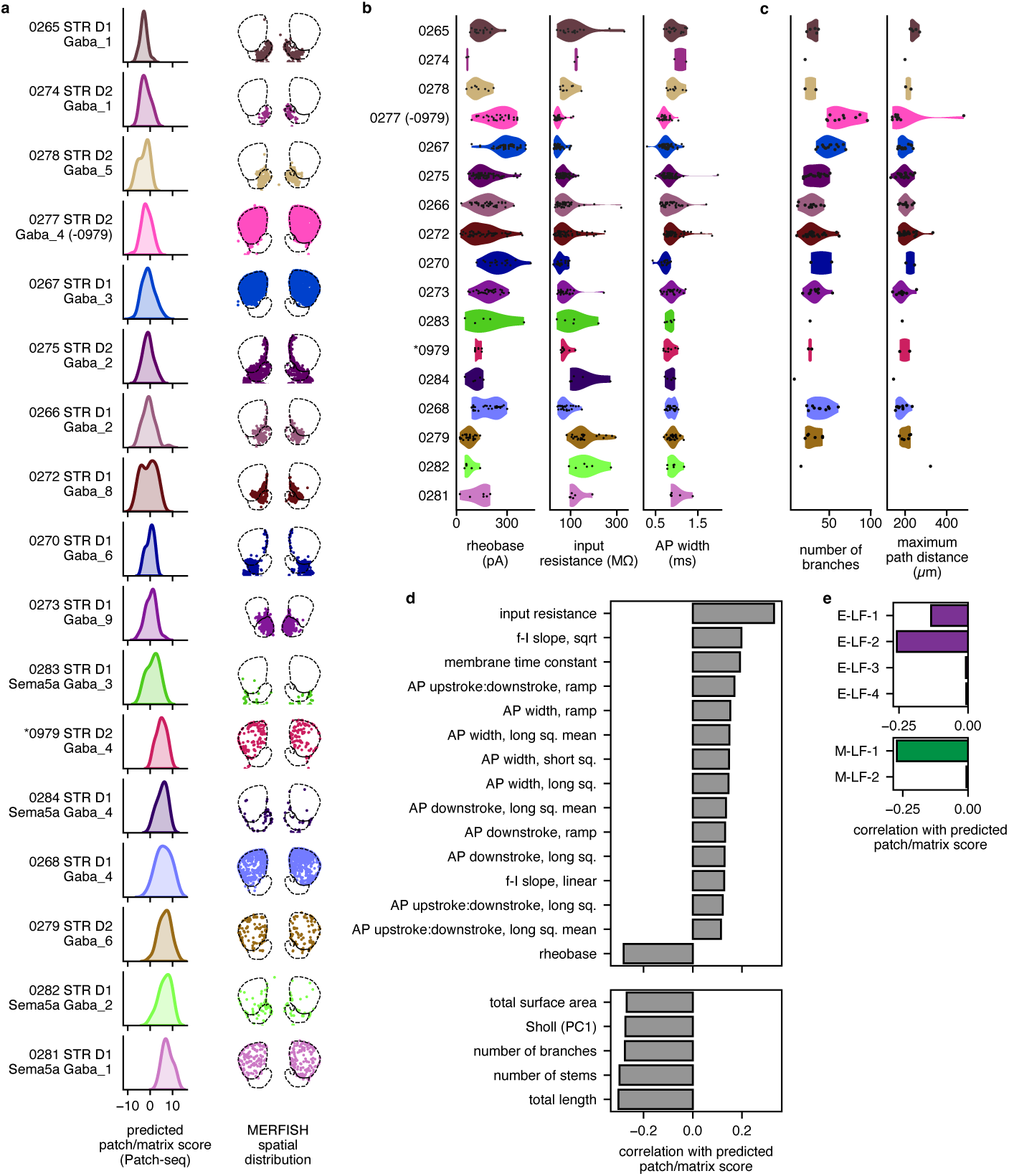
Patch/matrix score distribution and feature correlations. **a**, Distribution of patch/matrix score across Patch-seq cells mapping to MSN supertypes (left), and MERFISH spatial distribution for each supertype (right), ordered by predicted patch/matrix score median. T-type 0979 STR D2 Gaba 4 has been removed from the otherwise matrix-enriched supertype 0277 STR D2 Gaba 4, and shown in a separate row. Transcriptomic groups with a minimum of 6 cells are shown. **b**, Electrophysiological characteristics of Patch-seq cells mapping to MSN supertypes. As in **a** cells mapping to T-type 0979 STR D2 Gaba 4 have been removed from supertype 0277 STR D2 Gaba 4, and shown in a separate row (*0979 STR D2 Gaba 4). **c**, Morphological characteristics of MSN supertypes in **b**. **d**, Significant correlations between predicted patch/matrix score and electrophysiology (top) and morphology (bottom) features in Patch-seq cells. **d**, Correlations between predicted patch/matrix score and electrophysiology latent factors (top) and morphology latent factors (bottom) in MSNs. E-LF-1, E-LF-2 and M-LF-1 correlations were statistically significant.

**Extended Data Figure 8:**
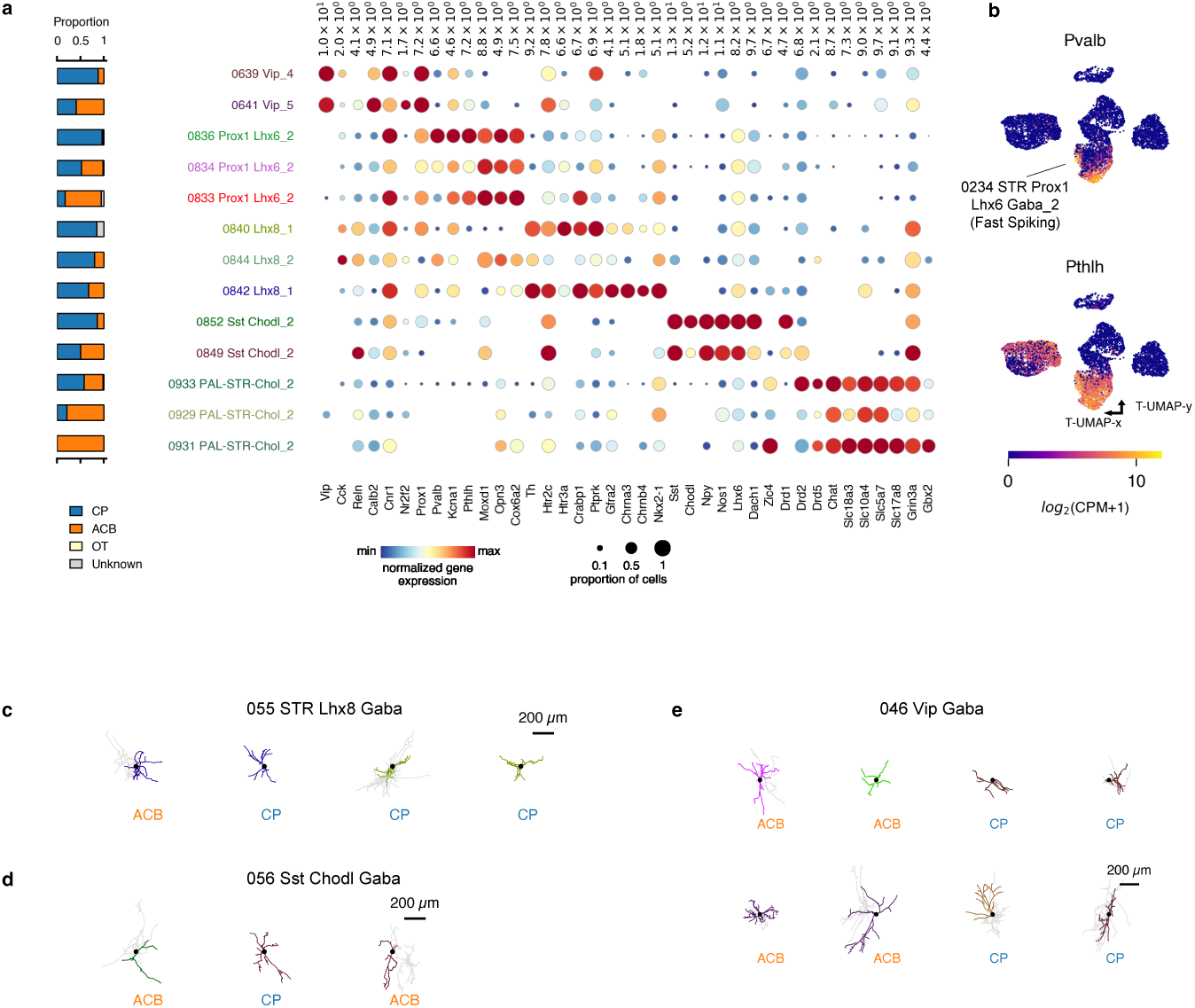
Properties of striatal interneurons. **a**, Summary of spatial localization and gene expression for striatal interneuron T-types: proportion of Patch-seq somas registered to CP/ACB/OT (left); select marker gene expression, normalized to maximum expression of each gene across interneurons (right). **b**, UMAP of gene expression of reference dissociated (n=6,225) and Patch-seq (n=395) striatal interneurons, colored by expression of *Pvalb* (top) and *Pthlh* (bottom). **c-e**, Example morphologies of neurons from the STR Lhx8, Sst Chodl, and Vip Gaba subclasses.

